# High-resolution diploid 3D genome reconstruction using Pore-C data

**DOI:** 10.1101/2023.08.29.555243

**Authors:** Ying Chen, Zhuo-Bin Lin, Shao-Kai Wang, Bo Wu, Long-Jian Niu, Jia-Yong Zhong, Yi-Meng Sun, Xin Bai, Luo-Ran Liu, Wei Xie, Ruibang Luo, Chunhui Hou, Feng Luo, Chuan-Le Xiao

**Author notes:** To whom correspondence should be addressed: Ruibang Luo.; Chunhui Hou.; Feng Luo.; Chuan-Le Xiao. These authors contributed equally to the manuscript as first authors (Ying Chen, Zhuo-Bin Lin, Shao-Kai Wang, Bo Wu, Long-Jian Niu, Jia-Yong Zhong).

## Abstract

In diploid organisms, spatial variations between homologous chromosomes are essential to many biological phenomena. Currently, it is still challenging to efficiently reconstruct a high-quality diploid 3D human genome. Here, we introduce Dip3D, reconstructing the diploid 3D human genome using Pore-C data of one sample. Dip3D has solved multiple problems in genome-wide SNV calling and haplo-tagging caused by the high sequencing error rates in Pore-C type data. Dip3D capitalizes on the high-order chromosomal interaction characteristics, enabling robust haplotype imputation and intricate haplotype-specific 3D structure discovery. Dip3D outperforms previous methods in data utilization rate, contact matrix resolution, and completeness by one order of magnitude. Moreover, Dip3D allows capturing haplotype high-order interactions that are unseen in Hi-C type data. We demonstrated the identified haplotype substructures such as Topologically Associating Domains (TADs) in the constructed 3D human genome, and unraveled connections between genic haplotype-specific high-order interactions and imbalanced allelic expression.

## Introduction

Chromosomes inside the cell nucleus are folded into three-dimensional (3D) structures featuring a multi-scale, hierarchical pattern encompassing A/B compartments, topologically associating domains (TADs), and chromatin loops^1^. This 3D genomic organization is central to numerous biological processes, including epigenetic regulation and gene expression^2,3^, DNA replication and repair^4,5^, and cell development and differentiation^5^, among others. In diploid organisms, spatial variations between homologous chromosomes are linked to allelic differences in gene expression^6^, DNA methylation^7^, and histone modification states^8^. They are also essential to biological phenomena such as female X-chromosome inactivation^9^ and early mammalian embryonic development^10^. Therefore, reconstructing the diploid 3D genome structure is critical for understanding a wide range of vital biological processes.

Combining chromatin conformation capture technologies (3C) and high-throughput sequencing technologies offers a promising approach to accurately delineating the diploid 3D genome structure. Previous diploid 3D genome structure reconstruction methods, such as HARP^11^, HiC-Pro^12^, and HaploHic^13^, were built upon Hi-C^14^, a technique that merges 3C with next-generation sequencing (NGS). However, the short read lengths and GC bias of NGS data have limited the proportion of haplotype-assigned Hi-C contacts. For example, there are only 0.14%-0.26% pre-imputation^3^,^13^ and ∼5.8% post-imputation^11,12^ Hi-C contacts with both ends being assigned to haplotype in the human cell line HG001, which leads to the requirement of a significant amount of data for reconstructing a diploid human 3D genome. In addition, the short fragment length makes it difficult to cover genomic regions with low phased single nucleotide variant (SNV) density, repetitive sequences, or high/low GC contents. Even with massive Hi-C sequencing data (over Tb), the completeness and continuity of reconstructed diploid human 3D genome structures remain limited^3,12^. Moreover, Hi-C unravels 2-way interactions, while realistic 3D genome structures involve concurrent high-order chromatin interactions in space, potentially with biological functions ^15^. NGS-based high-order chromatin interaction detection methods like Split-Pool Recognition of Interactions by Tag Extension (SPRITE)^16^ could hardly reveal higher than 3-way chromatin interactions. And reconstructing a superb high-order diploid human 3D genome remains a big challenge^17^.

Recently, researchers have developed Pore-C by integrating 3C with Nanopore long-read sequencing, which generates contact fragments (∼600-1000 bp) considerably longer than Hi-C reads (usually ≤150 bp)^3,18,19^. On this basis, we developed high-throughput Pore-C through the optimization of the library construction procedure and improved the sequencing yield by ∼80% on the same sequencing platform without changing the characteristics of the reads from ordinary Pore-C^20^. Pore-C can profile over 10-way chromatin interactions, unveiling synergistic high-order 3D chromatin interactions among regulatory elements^15,20,21^. Furthermore, unlike Hi-C, which relies on whole-genome phased SNVs from other technologies^17^, the long reads and abundant pairwise contacts of Pore-C enable independent genome-wide SNV detection and phasing, potentially allowing the reconstruction of diploid 3D genome structure using a single sample sequenced by one technology.

However, reconstructing diploid 3D genome structure from solely Pore-C data still faces multiple challenges. First, Pore-C inherits the high sequencing error rates and uneven error distribution associated with nanopore sequencing^22^, as well as sequencing errors tied to amino acid residues from the cross-linking phase during the Pore-C library preparation ^20^. These factors complicate SNV identification and haplo-tagging for Pore-C reads. The high sequencing errors and high-order contacts also lead to more h-trans (inter-haplotype) contacts in the Pore-C reads. Filtering out h-trans contacts and maximizing accurate h-cis (intra-haplotype) contacts become more complex for Pore-C than for 2-way Hi-C contacts. Second, even with the extended fragment lengths, only a few contacts (∼6.1%) can be haplo-tagged in human Pore-C data. Haplotype imputation is also essential for using Pore-C to construct accurate and high-resolution diploid 3D genome structures. However, while it is known that ∼99% of Hi-C contacts are h-cis contacts^23^, whether the high-order Pore-C reads dominantly contain fragments from the same chromosomes remains unclear. Hence, deciphering the haplotype constitution characteristics of high-order reads is fundamental for developing effective haplotype imputation strategies.

Here, we developed Dip3D to construct diploid 3D genome using Pore-C data of one sample. For SNV calling, we retrained a deep learning model for Pore-C^24^, which achieved over 95% recall and precision on ≥30× Pore-C data. Then, we developed a Pore-C-based SNV phasing pipeline and a Pore-C read haplo-tagging module, which reduced the overall h-trans rate from 10.1% to 5.6% while only dumping 4.10% haplo-tagging candidate fragments. To study the haplotype constitution of Pore-C reads, we generated F1 hybrid mice that had ∼3 times more heterozygous SNVs than HG001 and confirmed that the majority (>93%) of Pore-C reads are h-cis reads comprising fragments from the same haplotypes. Accordingly, we developed a progressive haplotype imputation strategy, improving the haplotype-assigned contact rate by 11.9-fold to 72.7% on human HG001 data. The diploid 3D contact matrix generated by Dip3D using less than half depth Pore-C (190×) data compared to Hi-C (531×) based matrix has achieved 6.9-fold haplotype-assigned contact density (1.80 vs 0.26 M/Gb), 10-fold higher resolution (5 kb vs 50 kb), and 63.9% fewer gap regions (9.35% vs 73.25%). Overall, Dip3D enables the unraveling of haplotype synergistic high-order 3D chromatin interactions using only a single sequencing technology and data from one individual sample.

## Results

### The design of Dip3D

The long fragment length and high-order interactions of Pore-C data make it superior for the reconstruction of diploid 3D genomic structure. To decipher diploid 3D genomes using Pore-C data, we developed Dip3D. The entire process of Dip3D is composed of seven steps (Figure 1): read mapping (1), genome-wide SNV calling (2), phasing (3), fragment haplo-tagging (4), haplotype imputation for untagged fragments (5), formatted output (6), and utilization of the high order diploid 3D genome map using supplementary tools (7). To develop Dip3D, we have also sequenced high depths of Pore-C data for human cell lines HG001 and HG002 and F1 hybrid mice in this study (Supplementary Table 1 and Supplementary Note 1).

In the first step, we utilize Falign, a tool specifically designed for fragmented long noisy reads^25^, to map Pore-C reads to the reference genome. Falign generates a specially formatted SAM output that records both the alignments and the arrangements of fragments in the high-order reads. For SNV calling using aligned Pore-C reads, we implement the new Pore-C model of Clair3 trained in this study, which has achieved high accuracy and recall rate.

In the SNV phasing step, Dip3D converts filtered high-order Pore-C reads into pairwise contacts and feeds them together with the called SNVs to HapCUT2^26^ for phasing, which achieves genome-scale high quality phased SNVs on human data. Then Dip3D uses the specifically developed Pore-C haplo-tagging module to proceed haplo-tagging of Pore-C fragments. The module has implemented three filters related to mapping quality (MAPQ), alignment identity, and SNV context mismatch count to significantly reduce the h-trans rate. After haplo-tagging, Dip3D performs haplotype imputation for untagged Pore-C fragments, leading to over ten-fold improvement in the data utilization rate. The haplotype imputation module primarily implements three progressive steps based on the haplotype constitution characteristics of high-order Pore-C reads, which were learned from the sequenced F1 hybrid (C57BL/6J × PWK/PhJ) mice and human Pore-C data.

Finally, Dip3D splits haplotype-assigned fragments of Pore-C reads into h-cis fragment groups saved in specialized SAM format (see Methods), from which diploid contact matrices (like Hi-C output) and haplotype high order interactions can be conveniently extracted using our supplementary tools (steps 6 and 7 in Figure 1). In this study, we also performed evaluation of the diploid 3D contact matrixes generated by Dip3D and leveraged the diploid highorder interactions to uncover new insights into biological phenomena.

**Figure 1.**
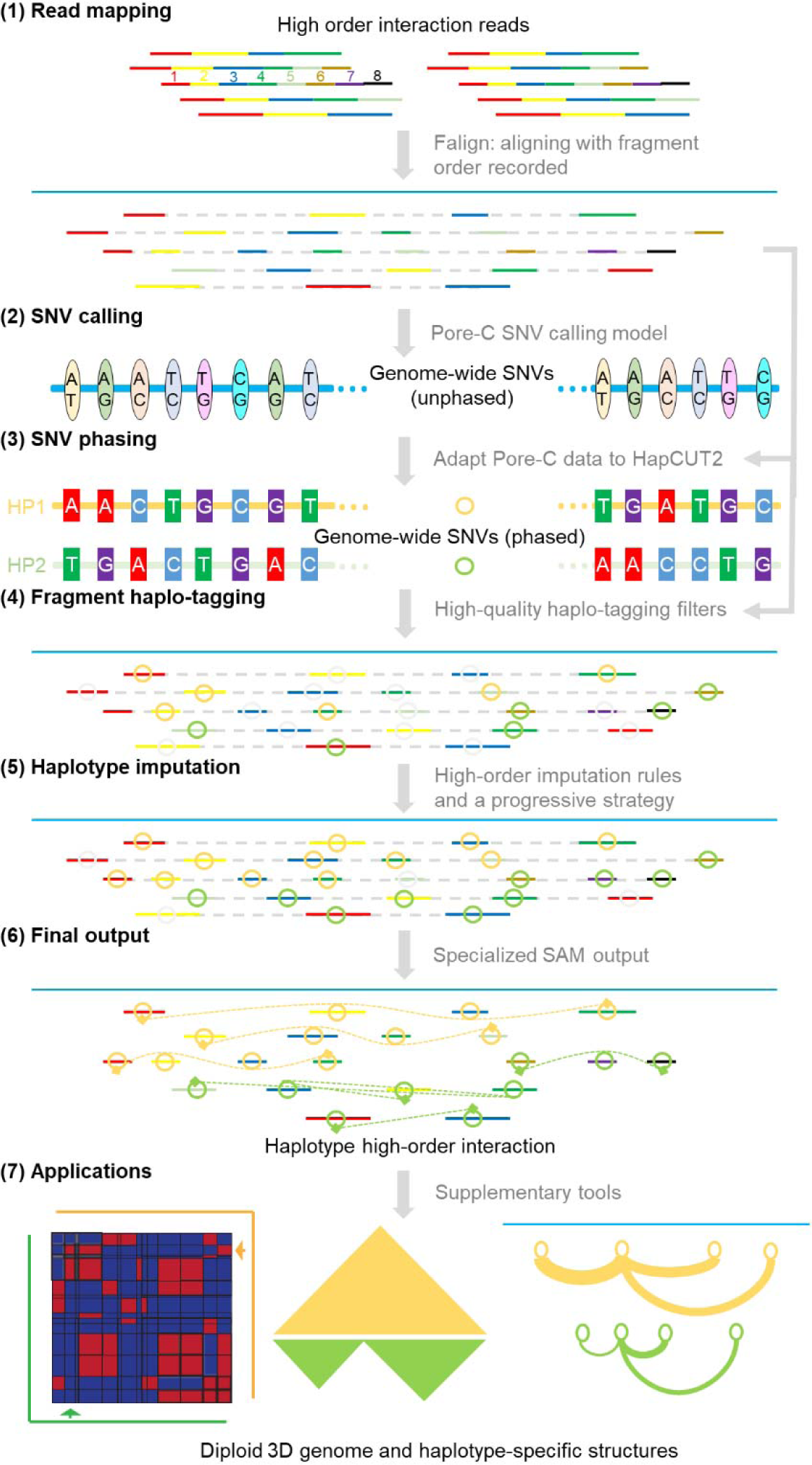
Schematic of Dip3D for constructing diploid 3D genomes. (1) Pore-C concatemers are decomposed and their fragments, whose orders in reads are represented by various colors, are mapped to the reference genome using Falign. (2) Genome-wide SNVs are identified from aligned Pore-C reads employing a specific Pore-C model for Clair3 trained in this study. (3) HapCUT2-based pipeline is used for phasing of the genome-wide SNVs, utilizing Pore-C reads. (4) Noisy Pore-C fragments are haplo-tagged using three optimized filters, identified through statistical analysis, with fragments assigned to haplotype 1 (HP1) and 2 (HP2) denoted by orange and green circles, respectively. (5) Haplotype imputation is carried out for untagged fragments (gray circles) in high-order reads, guided by the three rules learned in this study and implemented in a stepwise manner. (6) The final output Dip3D contains h-cis fragment groups and the fragment orders. Interaction paths are depicted as dashed curves linking fragments according to their orders in the original reads with two terminal fragments shown as dash squares. (7) Exemplary applications of the high-order diploid 3D genome: left, diploid contact matrices; center, intricate haplotype-specific 3D structures; right, haplotype-specific high order interactions.

### Training a Pore-C SNV calling model

Detecting heterozygous SNVs with high accuracy and recall rate is essential for high-quality diploid 3D genome reconstruction. However, compared to the corresponding benchmark SNVs from Genome in a Bottle (GIAB) in high confidence regions, the original Clair3 model (r941_prom_hac_g360+g422) only achieved recall rates of 69.8% and 87.6%, and precision rates of 72.8% and 88.2% on tested HG001 (30×) and HG002 Pore-C data (30×), respectively (Table 1). We then retrained a Clair3 model for Pore-C data using different depths of HG002 data sequenced in this study (see Methods). After training, the Pore-C Clair3 model reached SNV recall rates of 95.5% and 95.4%, accuracies of 98.5% and 95.2%, and F1 scores of 97.0% and 95.3% on the same HG002 (30×) and HG001 (30×) datasets, respectively (Table 1). With increases in the depth of Pore-C data to 60× and 90×, the Pore-C model of Dip3D recalled 97.9% and 98.6% heterozygous SNVs with ≥99% precision in the HG001 genome, achieving over 27.1% and 18.9% improvements of recall and precision rates compared to the original Clair3 model at the same sequencing depths, respectively. Importantly, Dip3D recalled ∼1.76-fold SNVs using Pore-C data in NGS-‘difficult’ regions compared to Hi-C based method at the same 90× depth (Supplementary Table 2). In summary, the high accuracy and recall rate of the new Pore-C model should make it suitable to call genome-wide SNVs for reconstructing high-quality diploid 3D genomes.

**Table 1.** Performance of Clair3 models on SNV calling from Pore-C data.

| Depth | Data type | Model | True positive SNVs | Recall | Precision | F1 score |
| --- | --- | --- | --- | --- | --- | --- |
| HG001 |  |  |  |  |  |  |
| 30x | ONT<sup>#</sup> | original | 2,873,218 | 0.997 | 0.996 | 0.996 |
|  | Pore-C | original | 2,011,117 | 0.698 | 0.728 | 0.712 |
|  | Pore-C | Pore-C | **2,752,867** | **0.955** | **0.985** | **0.970** |
| 60x | ONT | original | 2,879,669 | 0.999 | 0.997 | 0.998 |
|  | Pore-C | original | 2,041,529 | 0.708 | 0.790 | 0.747 |
|  | Pore-C | Pore-C | **2,822,855** | **0.979** | **0.996** | **0.987** |
| 90x | ONT | original | 2,880,419 | 0.999 | 0.997 | 0.998 |
|  | Pore-C | original | 2,049,498 | 0.711 | 0.808 | 0.756 |
|  | Pore-C | Pore-C | **2,844,219** | **0.986** | **0.997** | **0.991** |
| HG002 |  |  |  |  |  |  |
| 30x | ONT | original | 2,828,203 | 0.997 | 0.995 | 0.996 |
|  | Pore-C | original | 2,484,887 | 0.876 | 0.882 | 0.879 |
|  | Pore-C | Pore-C | **2,706,817** | **0.954** | **0.952** | **0.953** |
| 60x | ONT | original | 2,834,333 | 0.999 | 0.994 | 0.997 |
|  | Pore-C | original | 2,612,764 | 0.921 | 0.937 | 0.929 |
|  | Pore-C | Pore-C | **2,757,999** | **0.972** | **0.974** | **0.973** |
| 90x | ONT | original | 2,834,841 | 0.999 | 0.994 | 0.997 |
|  | Pore-C | original | 2,663,828 | 0.939 | 0.950 | 0.945 |
|  | Pore-C | Pore-C | **2,768,523** | **0.976** | **0.979** | **0.978** |

Note: <sup>#</sup>ONT means genomic nanopore sequencing data.

### Genome-wide SNV phasing and haplo-tagging methods

After the SNV calling, we developed a SNV phasing pipeline for Pore-C data. First, the pipeline converts multi-way contacts from Pore-C reads into pairwise contacts. Then, non-concatemer Pore-C reads (5-15% of total reads and can be seen as ordinary genomic reads) and pairwise contacts whose both fragments overlap with heterozygous SNPs were extracted and converted to compatible format for SNV phasing using HapCut2 (see Methods). The results of our pipeline on 30× HG001 Pore-C data already approached saturation on SNV phasing rate (Figure 2A) and achieved a phased SNV density of 0.70/kb, which is 1.72 and 1.18 times of those obtained using 30× and 531× Hi-C (Supplementary Table 3). Moreover, our pipeline obtained chromosome-level phased blocks and high phasing accuracy (Supplementary Table 3), with low switch and hamming error rates in both HG001 (0.59% and 0.44%) (Figure 2B,C) and the F1 mice (0.04% and 0.03%) data (Supplementary Figure 1A).

Next, we employed the phased SNVs to haplo-tag the Pore-C fragments. Fragments that contain phased SNVs are candidates for haplo-tagging. Roughly 21.3% of HG001 Pore-C fragments can be used for haplo-tagging on chromosome 1 (Figure 2D), which is 7.1 times the percentage observed in HG001 Hi-C fragments (3.0%). The percentages of haplo-tagged fragments increase with the length of the captured fragments(Figure 2E). On average, a Pore-C read contains 5.3 fragments, and at sequencing depths of 30×-90×, the phased SNV density is 0.70-0.77/kb for HG001. Therefore, most HG001 Pore-C reads containing 1-2 fragments as haplo-tagging candidates typically overlap with only 1-2 phased SNVs. As a result, precise identification of SNV alleles for accurate haplo-tagging on Pore-C fragments is a challenge. Using Whatshap^27^ for haplo-tagging HG001 Pore-C reads obtained a markedly higher h-trans error rate for Pore-C pairwise contacts compared to that obtained from Hi-C at the same interaction distances (Figure 2F). The rate even surpasses 5% when the fragment distance is less than 1 Mb, implying that the high sequencing error rate of Pore-C reads drastically compromises the precision of haplo-tagging.

To achieve high-performance haplo-tagging, we implemented several optimizations in Dip3D. First, we set up the optimized thresholds of mapping quality (MAPQ) score and percentage of identity (PI) for haplo-tagging. By default, WhatsHap disregards alignments with a MAPQ score below 20 to increase accuracy, but this also leads to a loss of candidate Pore-C fragments for haplo-tagging. To recall more haplo-tagging candidates, we tested various MAPQ and PI threshold and determined MAPQ ≥ 5 and PI ≥ 85% as the optimal combination (Figure 2G, H). These parameters facilitated the recovery of high-confidence haplo-tagged fragments with MAPQ<20, while maintaining a relatively low h-trans rate (<5%). Second, considering the uneven distribution of sequencing errors and the prevalence of deletions in nanopore reads^22^, which affects local alignment accuracy and can lead to false haplo-tagging, we required that no mismatch (potential sequencing error) occurs in the fragment bases upstream and downstream of the SNV sites to be used for haplo-tagging. Results showed that as the length of mismatch-free contexts surrounding the phased SNVs was extended, the h-trans error rate decreased accordingly (Figure 2I). Our strategy filtered out 4.1% of candidate fragments as untrustworthy, haplo-tagged about 20.4% of Pore-c fragments, and reduced the overall h-trans contact rate from 10.1% to 5.6% (Figure 2J), which is critical for the accurate construction of diploid 3D genome structures.

**Figure 2.**
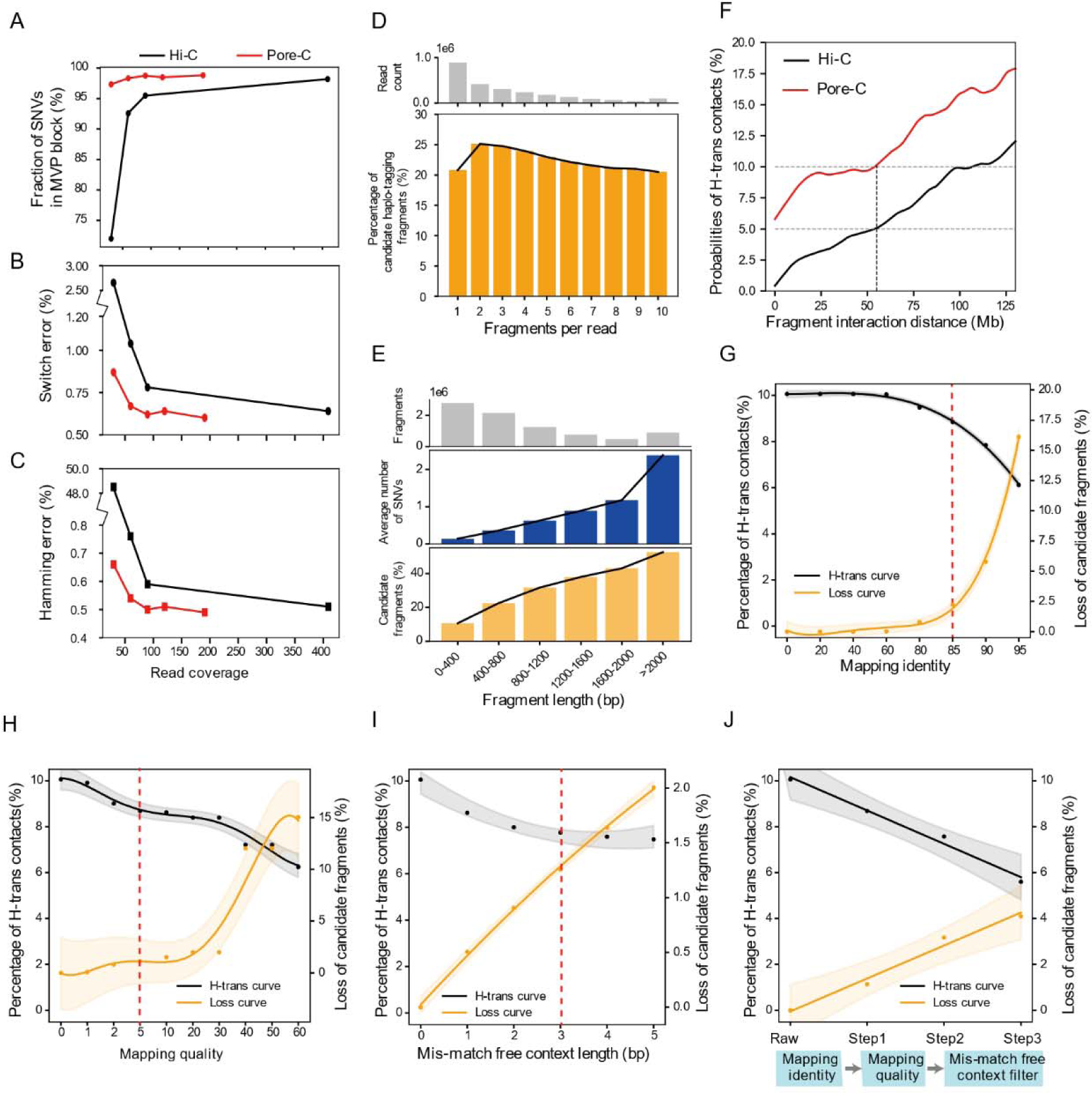
Statistics on SNV phasing and haplo-tagging. HG001 data mapped to the reference chromosome 1 are summarized in this figure. **A**-**C**, comparison of phasing completeness (A), switch error rate (B), and hamming error rate (C) between Hi-C and Pore-C at different sequencing depths. MVP block denotes the most-variants-phased block of a chromosome. **D** and **E**, distribution of candidate haplo-tagging fragment rates in Pore-C reads containing different numbers of fragments (D) and among Pore-C fragments of different lengths (E). **F**, relationship between h-trans rate and interaction distance for Pore-C and Hi-C pairwise contacts. Haplo-tagging was carried out using WhatsHap with default settings for both the Pore-C (190×) and Hi-C (531×) data. **G**-**I**, h-trans rates and candidate haplo-tagging fragment loss rates under different thresholds of fragment mapping identities (G), mapping qualities (H), and mismatch-free context lengths (I). **J**, cumulative h-trans rate decrease and candidate fragment loss after applying three filters.

### Characteristics of haplotype high-order interactions in Pore-C data

Previous study showed that h-cis contacts are dominant in pairwise contacts from Hi-C type 3C data^23^. However, it remains uncertain whether the fragments in the high-order Pore-C reads mainly originate from the same chromosomes. To address this question, we generated F1 (C57BL/6J × PWK/PhJ) hybrid mice (Figure 3A) that had approximately 10-times heterozygous SNV density compared to HG001 and sequenced 54× Pore-C data. We detected 80.4% (16,210,490 / 20,156,985) of putative F1 heterozygous SNVs (homozygous SNVs between parental lineages from public database) using the Pore-C data, ∼15.4% higher than previously reported using Hi-C data on a similar F1 strain^28^, suggesting high data quality. Using the F1 mice dataset, we first explored fragment haplotype distributions in Pore-C reads. The much higher heterozygosity led to 62.4% of the mice Pore-C fragments to be directly assigned to a haplotype via SNVs (Figure 3B), 2.8 times compared to the HG001 dataset (22.5%). High-order reads containing three or more tagged fragments made up 42.0% of all mouse Pore-C reads. Within this subset, 93.0% of the reads were h-cis (single haplotype) reads, which maintain an approximately equal distribution between paternal and maternal haplotypes (Figure 3C). The percentage of h-cis reads decreased as the number of haplo-tagged fragments per read increased, yet it remained above 92% across all categories (Figure 3D). Within fully haplo-tagged high-order reads (19.0% of the total Pore-C reads), 95% were identified as h-cis reads with ∼1:1 paternal to maternal read ratio (Figure 3E). Considering the h-trans errors due to false haplo-tagging, the proportion of h-cis Pore-C reads most likely has been underestimated. In summary, our findings verify that most Pore-C reads are indeed h-cis reads, and haplotype imputation among high-order interactions is theoretically feasible.

**Figure 3.**
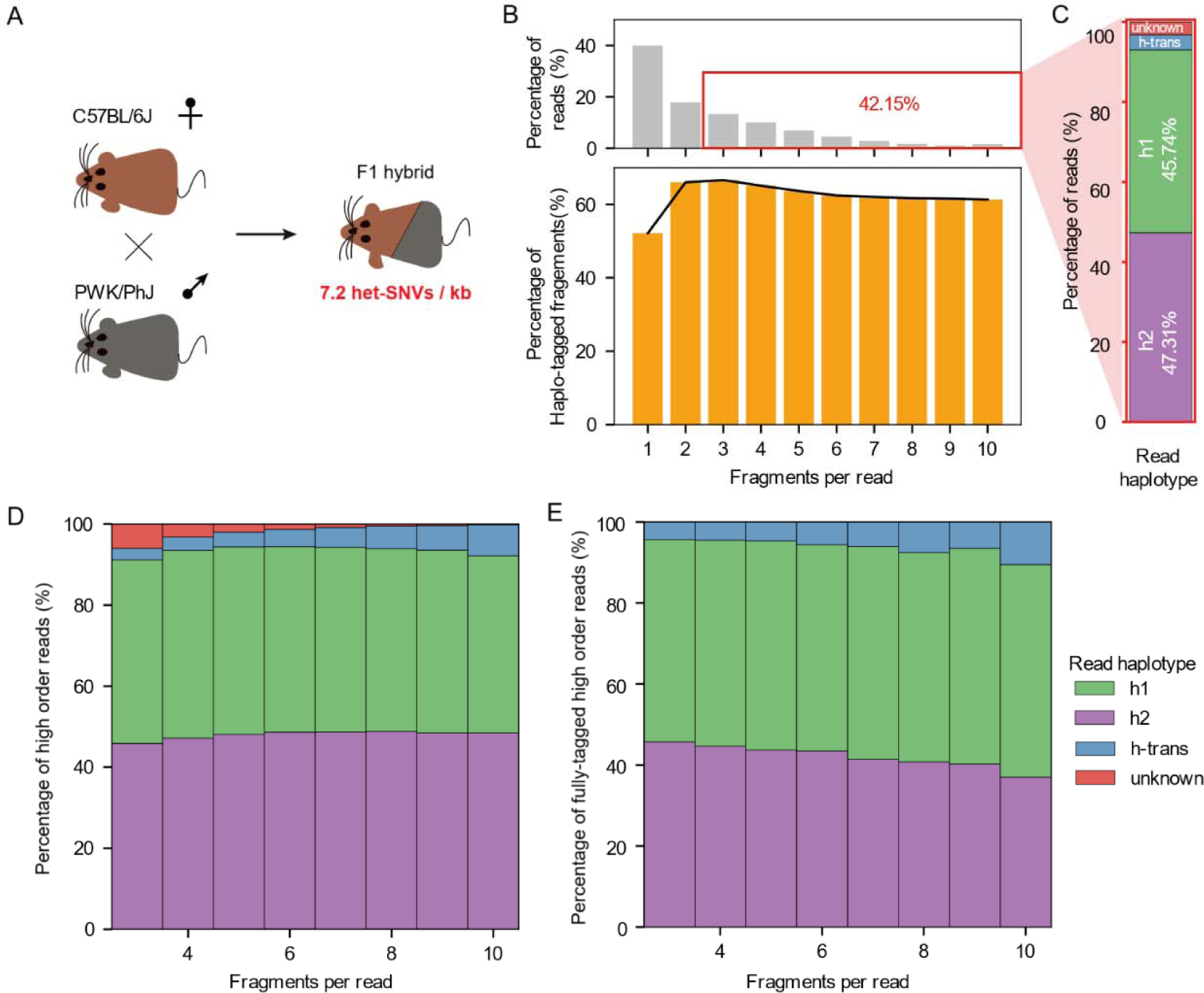
Haplotype composition of Pore-C reads from F1 hybrid mice. Graphs in this figure were based on the sequenced 54× Pore-C data of the F1 mice. **A,** generation of highly heterozygous F1 mice (C57BL/6J × PWK/PhJ). **B,** distribution of read fragment counts and fragment haplo-tagging ratios in the analyzed Pore-C dataset. **C**, haplotype distribution of high order (≥ 3-way) reads. Read haplotype categories: h1, reads containing maternal haplotype (and unassigned) fragments only; h2, reads containing paternal haplotype (and unassigned) fragments only; h-trans, reads containing both maternal and paternal haplotype (and unassigned) fragments; unknown, reads containing unassigned fragments only. **D** and **E**, haplotype distribution of all high-order reads (D) and fully haplo-tagged high-order reads (E) including different numbers of fragments.

### Haplotype imputation for Pore-C high-order contacts

While Pore-C has much higher (44.8-fold based on HG001 data) contact haplo-tagging rate than Hi-C, they only represent a small fraction (6.1%) in the human dataset (Supplementary Table 4). Therefore, it is critical to perform haplotype imputation to maximize correct h-cis contacts while limiting the h-trans error rate. Based on the characteristics of Pore-C data, Dip3D performs three steps of haplotype imputation (Figure 4A-D).

First, similar to Hi-C data^23^, significant positive correlation between interaction distance and h-trans contact rate of Pore-C contacts was observed in Pore-C data (Figure 4A). To limit false imputation rate, we selected the interaction distance (18 Mb) of Pore-C pairwise contacts with 5% h-trans rate as threshold for distinguishing short-range (≤18 Mb, reliable for imputation) and long-range (>18 Mb, less reliable) contacts. Moreover, adjacent (directly neighboring) contacts (20 Mb) had a larger interaction distance at 5% h-trans rate than the total pairwise contacts and therefore acquired a more relaxed 20 Mb distance threshold (Figure 4A). For each untagged fragment, we inferred its haplotype using its short-range contacts (Figure 4D). If an untagged fragment obtains a unique haplotype or it has at least two-fold short-range contacts with one haplotype compared to the other, it obtains the unique or the large-score haplotype identity, which is referred to as the distance rule of haplotype imputation.

**Figure 4.**
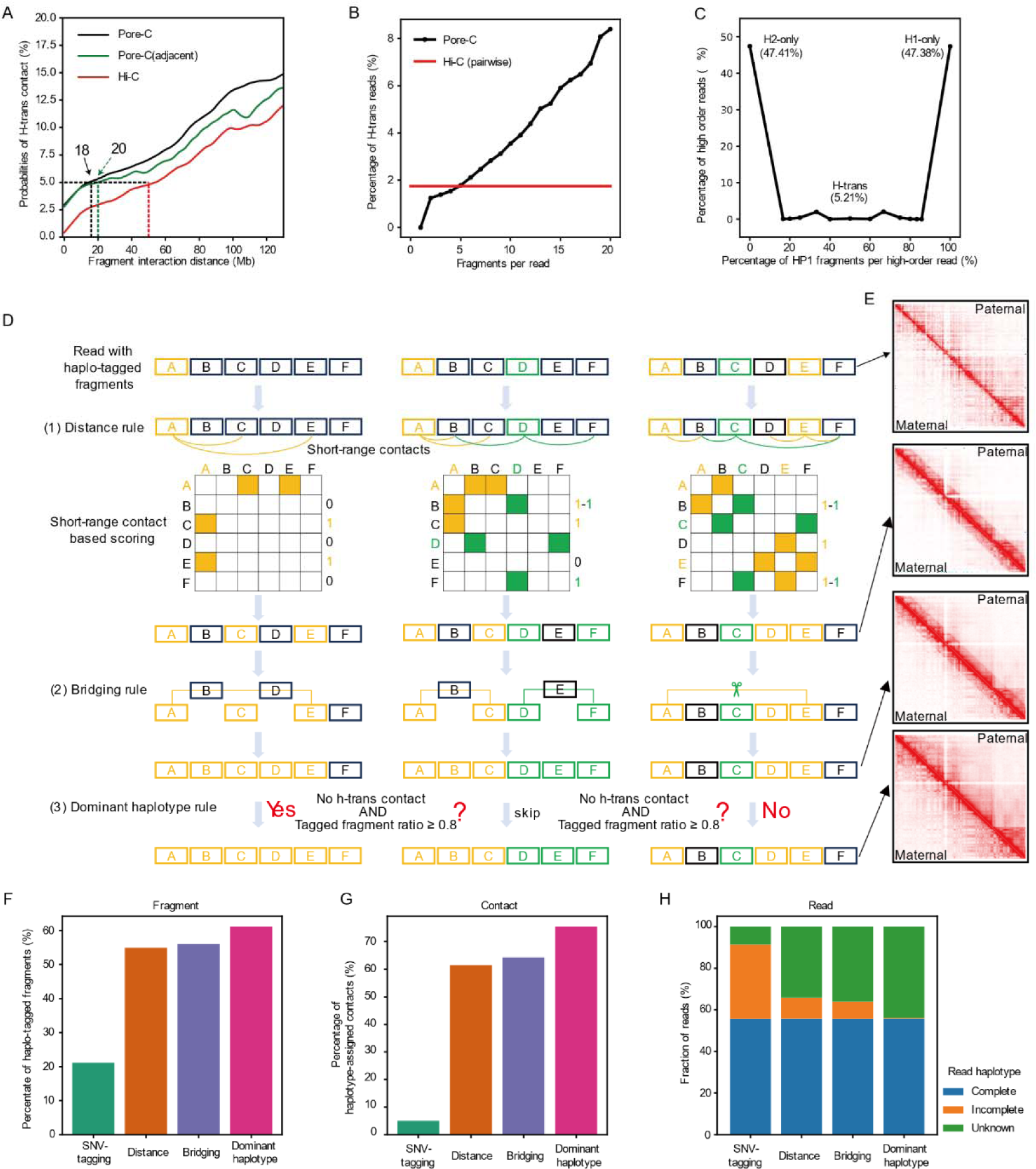
Haplotype imputation for high order Pore-C reads. **A**, correlation between interaction distance and h-trans rate. ‘Pore-C’ encompasses all pairwise contacts from Pore-C reads, and ‘Pore-C (adjacent)’ refers to pairwise contacts between directly neighboring fragments in Pore-C reads. Interaction distance thresholds for distinguishing short-range and long-range contacts are set at the 5% h-trans contact rate level. **B**, relationship between read fragment count and h-trans read rate. **C**, distribution of h-cis and h-trans high-order reads. **D**, execution of distance, bridging, and dominant haplotype rules in haplotype imputation for high-order reads. Rectangles A-F denote six fragments of a Pore-C read, and elements belonging to the two different haplotypes are colored in orange and green, respectively. **E**, diploid contact heatmaps of HG001 before and after each of the imputation steps. The arrows denote corresponding stages between the diagram (D) and the diploid contact heatmaps (E). **F**-**H**, proportions of haplo-tagged fragments (F), haplotype-assigned contacts (G), and complete/incomplete haplotype-assigned reads (H) after each of the three imputation steps.

Second, we found that the occurrence of a different haplotype fragments between two identically haplo-tagged fragments was rare (1.1%) in the mice Pore-C reads. Accordingly, we introduced a bridging rule to infer the haplotype of an internally untagged fragment based on surrounding haplo-tagged fragments in concatemers. For contacts that link two fragments with a same haplotype (called a bridge), if no fragment between those two fragments is tagged with a different haplotype, we assigned the untagged fragments across the bridge chain to the haplotype of the terminal fragments (Figure 4D).

Finally, most (∼93%) human and F1 mice Pore-C reads are h-cis reads (Figure 3C-E and Figure 4B,C), which allows imputation for untagged fragments with long interaction distances with the rest fragments. In the F1 mice data, the ratio of fragments belonging to the secondary haplotype (occupying ≤ 50% fragments) of a h-trans read is limited (1.3%) among reads with ≥3 haplo-tagged fragments. Accordingly, we proposed the dominant haplotype rule that assigns untagged fragments to the dominant haplotypes occupying ≥80% fragments of a read. We applied the dominant haplotype rule after the two steps above, with an extra requirement that no h-trans is observed in the read.

To show the effects of the three haplotype imputation steps, Figure 4E plots the pairwise contact heatmaps for paternal and maternal X chromosomes of HG001 before and after each step. The paternal-specific bipartite structure with two super-domains related to X chromosome inactivation became more easily visible after each imputation step. As shown in Figure 4E, the distance rule primarily recovered proximal contacts near to the diagonals, while the bridging and dominant haplotype imputation steps recovered more distal contacts further away from the diagonals. Before imputation, the ratio of haplo-tagged fragment was 22.5%, which was increased by 32.9%, 1.3%, and 4.6% after the three imputation steps respectively to a final 60.0% (Figure 4F and Supplementary Table 4). Correspondingly, the ratio of haplotype-assigned contacts was increased from 6.1% to 72.7%, with 56.4%, 3.6%, and 6.6% improvement after each step (Figure 4G). The percentage of fully haplo-tagged reads also increased from 7.7% to 42.4% after haplotype imputation (Figure 4H). These results indicate that our imputation strategy has effectively recovered proximal and distal contacts for high-quality diploid 3D genome structure construction.

### Advantages of diploid 3D genome at the pairwise contact level

For quality assessment, we extracted diploid pairwise contact matrices from the Dip3D output based on 190× HG001 data. We compared the diploid contact matrices with those based on 531× Hi-C data of Hg001 using the restriction endonuclease MboI, isoschizomer of DpnII used in this study, from the study of Rao et al.^3^. In summary, Dip3D matrices had 6.9-fold haplotype-assigned contact density (1.80 M/Gb vs 0.26 M/Gb), 63.9% fewer gap regions (9.4% vs 73.3%), and 10-fold resolution improvement (5kb vs 50kb, Figure 5A), which denotes the minimum bin size that ensures an overlap of ≥1,000 contacts in at least 80% of bins. To evaluate the continuity of diploid 3D structures, we proposed the concept of ungapped haplotype contact block, which denote continuous regions covered by haplotype-assigned 3C contacts. Due to the much longer fragment length and higher contact utilization rate, the Dip3D matrices achieved a remarkable 5,000-fold ungapped haplotype contact block N50 of 14.5 Mb compared to Hi-C matrices (2.9 kb). We also assessed the ability of the HG001 Dip3D matrices in recovering canonical haplotype-specific 3D structures previously revealed by high-depth (878×) Hi-C^3^. With no surprise, haplotype-specific structures related to X-chromosome inactivation (XCI) and genetic imprinting are easily visualized in Dip3D matrices (Supplementary Figure 2 and Supplementary Note 9).

We further compared the coverage of the Pore-C (190×) and Hi-C (531×) based diploid contact matrices of HG001 across different types of NGS ‘difficult’ regions. The results showed that Dip3D matrices had near-complete (≤2.1% gaps) coverage in high/low GC content regions, whereas the Hi-C matrices had 69.5-97.2% gaps (Figure 5B and Supplementary Table 5). In different types of repeats regions, the Dip3D matrices also had an average of 59.8% lower gap ratios than the Hi-C matrices had (Figure 5B). The Hi-C matrices exhibited significantly lower contact frequencies in all types of ‘difficult’ regions compared to non-‘difficult’ regions, whereas the Dip3D matrices only showed significantly lower contact frequencies in segment duplication and low-mappability regions (Figure 5C). For each type of ‘difficult’ region, the frequency of haplotype-assigned contacts was significantly higher in the Dip3D matrices than in the Hi-C diploid contact matrices (Figure 5C).

Pore-C based Dip3D matrices could have advantages over the Hi-C based matrices even with the same contact size. To confirm this, we compared randomly down sampled HG001 Dip3D matrices (160M contacts per haplotype, equaling 59× Pore-C data) to the equal contact sized diploid contact matrices from 878× Hi-C data^3^. Both matrices achieved 50kb resolution. However, the Dip3D matrices displayed a more even contact frequency distribution than the Hi-C matrices did, which showed considerably higher variances and interquartile ranges, and an obvious right-skew (Figure 5D). These results indicate that the genomic distribution of haplotype contacts is much less biased in the Dip3D matrices than in the Hi-C based matrices.

Then we compared the noise signal levels of the same equal sized Dip3D and Hi-C matrices using the QuASAR-QC score, which estimates the quality of 3C type data at different resolutions using cis-interaction correlations among spatially close bins^29,30^. The results showed that the HG001 Dip3D matrices had higher QuASAR-QC scores than the Hi-C diploid matrices at all resolutions (Figure 5E and Supplementary Table 6). The qualities of the Hi-C diploid matrices also declined more rapidly than the Dip3D matrices as the resolution increased (i.e., bin size reduced). These findings also suggest that Dip3D has effectively mitigated the impact of Pore-C’s noise signals on the final diploid 3D genomic structure.

The more even genomic distribution of the Pore-C diploid contacts might help discover 3D structures previously undetected by Hi-C. Indeed, the Pore-C matrices (160M × 2) reproduced 2.3 times more TAD structures (unphased, detected using 878× Hi-C) than those detected from the 160M × 2 Hi-C matrices (Figure 5F and Supplementary Table 7). Most of the additional TADs were related to NGS ‘difficult’ or SNV-sparse regions that had sparse Hi-C contact coverage. For instance, the Hi-C matrices had sparse haplotype-assigned contacts in an exemplary region (blue) on chromosome 1, making the TAD structures unclear, while the TAD structures were clearly discernible in the Pore-C matrices (Figure 5G). Another instance was the chrX:47.5-49.5 Mb region, in which the Hi-C matrices were almost blank, while different contact frequencies could be observed between the maternal and paternal haplotypes in the Pore-C matrices (Supplementary Figure 3). Surrounding the chrX:48.4-48.6Mb region, a prominent TAD structure was presented in the maternal haplotype but absent in the paternal haplotype. Our findings echo a previous study that identified five maternally-expressed genes in this region using RNA-seq and CIA-PET RNAPII^6^.

Overall, Pore-C-based diploid 3D genomic structure exhibits dramatic advantages in terms of contact density, resolution, completeness, and continuity (evidenced by ungapped haplotype contact blocks) compared to the Hi-C-based structure, and this superiority persists even when the Pore-C sequencing depth is significantly lower (59.2× vs 878×) than Hi-C.

**Figure 5.**
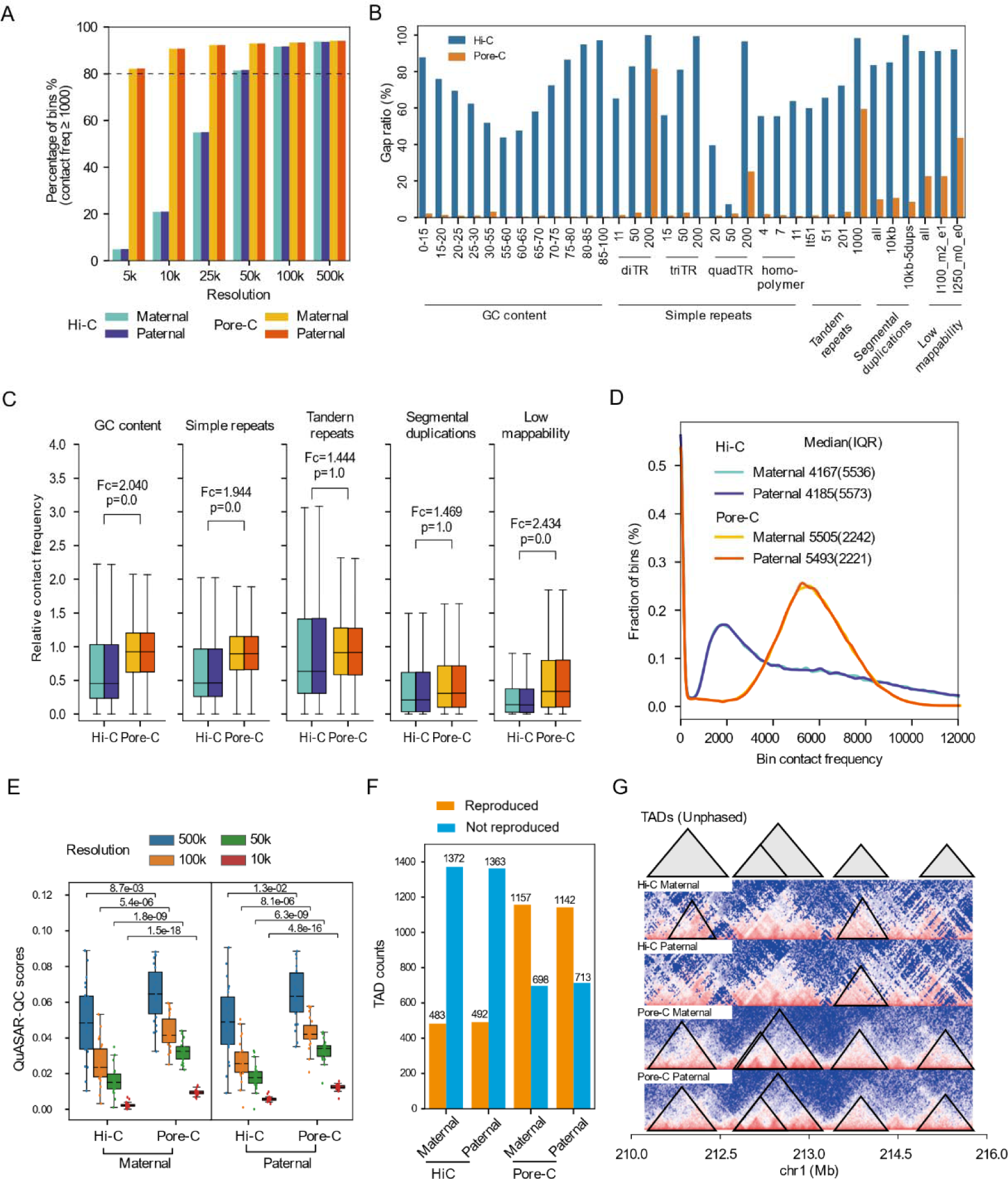
Evaluation of contact distribution and 3D structures in diploid 3D genomes. Panels **A** and **B** depict the results based on 531× Hi-C and 190× Pore-C data of HG001. **A**, resolution of Pore-C (Dip3D) and Hi-C based diploid 3D genomes. Resolution denotes the minimum bin size that ensures an overlap of ≥1,000 contacts in at least 80% (dash line) of bins. **B**, gap ratios of diploid contact matrices in NGS ‘difficult’ regions. In this graph, regions with <10 haplo-tagged fragments coverage were regarded as gap regions (Supplementary Table 5). Panels **C-G** compared the performances of equal contact sized (160M per haplotype) Pore-C (Dip3D) and Hi-C (Rao et al., 2014) diploid contact matrices. **C**, comparison of diploid contact frequency distributions in NGS ‘difficult’ regions. The relative contact frequency was normalized by dividing the mean haplotype-assigned contact frequency (1.0 on the axes) of the non-‘difficult’ regions. For both Hi-C and Pore-C, the maternal (left boxplots) and paternal (right boxplots) distributions were drawn separately. Fc denotes fold change. **D**, distribution of contact frequencies (50kb bins) in the diploid 3D genomes, comparing Pore-C and Hi-C. **E**, QuASAR-QC scores of Pore-C and Hi-C based diploid contact matrices at different bin sizes. Evaluation of the maternal (left) and paternal (right) haplotype matrices was carried out separately. **F**, statistics on the number of unphased Hi-C TADs reproduced in Hi-C and Pore-C based diploid 3D genomes. All TADs were called at 50kb resolution. Unphased TADs were called using the same Hi-C data used for diploid 3D genome reconstruction. Only TADs with reciprocal ≥ 80% overlap were regarded as being reproduced. A total of 1,855 unphased TADs (50kb resolution) across the genome were subjected to this test. **G**, TADs (unphased) reproduced from Hi-C and Pore-C diploid contact matrices in an exemplary region.

### Haplotype-specific high-order interaction and allele-specific expression

Allele-specific expression (ASE), also known as allelic expression imbalance, is potentially associated with haplotype-specific 3D genome structures, but the principles governing their association remain unclear. Dip3D enables investigation of haplotype-specific higher-order interactions and their links to ASE. Promoter high-order interaction analyses showed that the single chromatin distal (>3 Mb distance) to proximal (≤3 Mb) contact ratios (SDPR) at promoter regions were negatively correlated (*r*=-0.12, *p*<0.001) with gene expression (unphased), while the number of simultaneous interacting enhancers of promoters and multi-enhancer (ME) interacting read ratio were positively and more strongly correlated (*r*=0.45 and 0.41, *p*<0.001) with gene expression abundancy (Supplementary Figure 4A-E). To identify if they are also involved in ASE, we compared the SDPRs and enhancer interacting behaviors of promoters between gene alleles in HG001 Pore-C data. Since genetic imprinting on autosomes and X chromosome inactivation (XCI) are two different biological processes that both lead to ASE, we investigated them separately. For imprinted and unimprinted genes on the autosomes, we didn’t observe significant variation between them in allelic SDPR or allelic multi-enhancer interaction (Figure 6A,B and Supplementary Tables 8,9). There was no significant difference being observed between alleles of previously reported imprinted genes in respect of either SDPR or multi-enhancer interaction (Supplementary Tables 8,9).

Unlike genetic imprinting, haplotype specific high-order interactions could have played an essential role in the XCI process. The promoters of genes on the paternal-origin X chromosome had significantly higher SDPRs than those on the maternal-origin X chromosome in the female cell line HG001 (Figure 6C). X chromosome genes also had significantly (*p*<0.001) larger allelic SDPR difference than those imprinted/unimprinted autosome genes had. However, there was no significant difference between genes subjected to XCI and those escaping from XCI in respect of allelic SDPR levels (Supplementary Figure 4F), indicating allelic SDPR difference is not directly related to ASE on the X chromosomes. To the contrary, maternal genes subjected to XCI had significantly more simultaneous interacting enhancers (Supplementary Figure 4F) and a significantly higher proportion of multi-enhancer interacting (ME) reads than the corresponding paternal genes (Figure 6D and Supplementary Table 9). Interestingly, genes that escape from XCI didn’t show significant difference in allelic multi-enhancer interaction (Supplementary Figure 4F).

Here, we have proposed a chromatin higher-order interaction model for XCI related ASE. Taking the chrX maternal ASE gene RBM3 as an example, although there are similar numbers of haplotype-assigned multi-way (≥5 fragments) reads between the maternal (n=248) and paternal (n=209) haplotypes, the maternal chromatin promoter of RBM3 interacts more frequently with proximal regions than the distal regions, whereas it’s opposite for the paternal chromatin, leading to significantly lower maternal SDPRs (Figure 6E). Maternal ME reads are significantly enriched in promoter-enhancer interaction patterns by ∼3-fold (0.337 vs 0.123) compared to the paternal ME reads (Figure 6F). We observed a similar trend on other genes subjected to XCI and detected with ASE, which had significantly enriched ME reads (Supplementary Table 9). Accordingly, as shown in Figure 6G, the promoters of XCI-related genes have more intensified proximal interactions in the maternal chromosome, which also interact with more enhancers to form interaction hubs, while their paternal promoters interact more frequently with distal regions and with fewer regulatory elements. And such haplotype high-order interaction differences collaborate with other allelic epigenetic differences^31^ to cause ASE on female X chromosomes.

In summary, chromosome compaction difference (indicated by SDPR) and high-order regulatory element interactions both have played a role in XCI, but only haplotype-specific regulatory element interaction is directly involved in ASE related to XCI. On the contrary, genetic imprinting on the autosomes tend to be regulated by other mechanisms such as DNA methylation^32^ than haplotype-specific high-order interactions.

**Figure 6.**
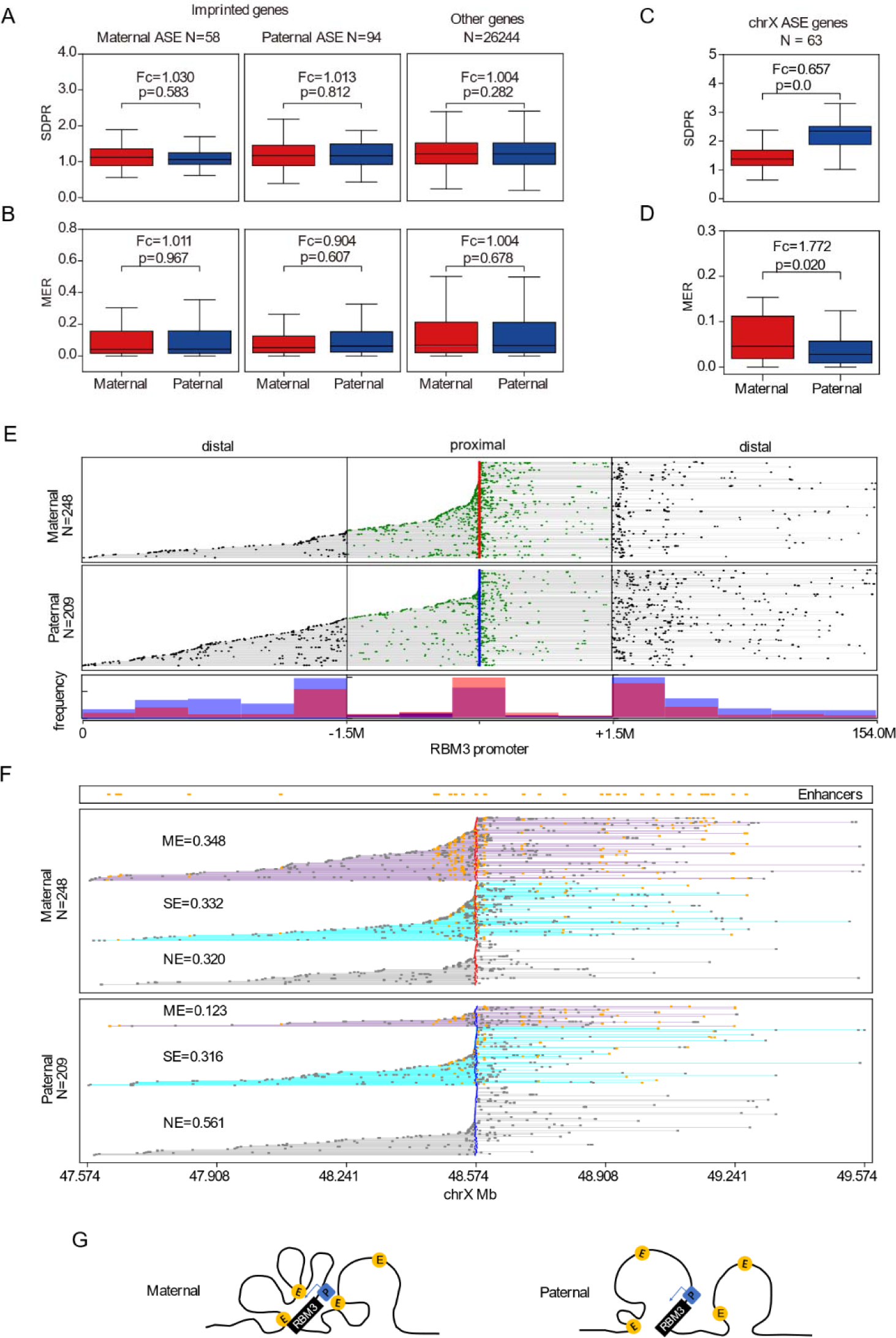
Haplotype-specific high-order interaction at genic promoter regions. **A** and **B,** distribution of allelic single chromatin distal to proximal contact ratios (SDPR) and multi-enhancer interacting ratios (MER) at the promoter regions of previously reported imprinted genes and other genes. Imprinted genes are classified as maternal (left) and paternal (right) ASE (allele-specific expression) according to which allele has been reported to be dominant in expression. **C** and **D**, distribution of allelic SDPRs and MERs at the promoter regions of previously reported ASE genes on the X chromosome of HG001. **E**, distribution of distal and proximal RBM3 promoter interacting fragments. In the top h1 and h2 panels, each row represents one high order read with differentially colored proximal (green dots) and distal (black dots) interacting fragments. **F**, distribution of reads with different RBM3 promoter-enhancer interacting patterns (NE, SE, and ME). NE, SE, and ME denote the promoter interacting with zero, a single, and multiple enhancer(s), respectively. **G**, haplotype-specific high order interaction model (using RBM3 as an example) for ASE genes subjected to the impact of X chromosome inactivation.

## Discussion

Dip3D performs the entire process of constructing diploid 3D genome structure using only Pore-C data of one sample. It has resolved multiple challenges posed by the noisy high-order Pore-C reads in the steps, including SNV calling, SNV phasing, haplo-tagging, and haplotype imputation. The haplo-tagging and haplotype imputation strategies applied in this study dramatically increased the data utilization rate of Pore-C without compromising on the 3D genome structure quality. In our evaluation, Dip3D outperforms previous Hi-C based approaches in data utilization rate, contact map completeness, and 3D contact matrix density by over a magnitude. Most importantly, Dip3D enables learning haplotype-specific high-order interactions which potentially have biological significance.

The capacity to decode and reconstruct the three-dimensional structure of diploid genomes is important for the understanding of chromatin spatial organizations and many biological processes. Conventional methods rely on Hi-C that needs chromosome-level phased SNVs derived from additional sequencing technologies^28,33^ and high sequencing data coverage^3^, and only produce diploid 3D genome with limited completeness and resolution. Pore-C reads have much longer read lengths and contain higher order interactions, which gives Pore-C a distinct edge in diploid 3D genome analysis and mitigates the limitations inherent in Hi-C data, such as a low ratio of reads overlapping heterozygous SNVs, short read length, and GC bias. Using the hybrid F1 mouse, we demonstrated in this study that the majority of high-order Pore-C reads contain fragments from the same chromosomes, similar to that observed in the low-order Hi-C data^23^. This finding forms the foundation for developing haplotype imputation rules for Pore-C reads to achieve high resolution diploid 3D genome. The high-resolution diploid 3D genome allows researchers to identify more fine 3D genome structure and further understand the biological processes that are associated with the 3D structure of genome. Given the substantial improvements in accuracy and throughput of nanopore sequencing technology, Pore-C is increasingly utilized in 3D genome studies. Dip3D, with its distinct advantages, may become an essential tool in diploid 3D genome structure analysis.

## Methods

### Culture of human cell lines

Human B lymphocyte cell lines HG001 (NA12878; Coriell Institute, RRID:CVCL_7526) and HG002 (NA24385; Coriell Institute, RRID:CVCL_7526) were cultured in 1× RPMI 1640 medium, which was supplemented with 15% fetal bovine serum (FBS) and 1% penicillin/streptomycin. The culture conditions were maintained at 37°C with 5% CO_2_. The cells were passaged twice a week or whenever the cultures achieved 80% confluence.

### Obtaining F1 hybrid mice by in vitro fertilization

Hybrid mice were generated using in vitro fertilization. The female C57BL/6J mice, aged 5-6 weeks, were first treated with 5 IU of pregnant mare’s serum gonadotropin (PMSG) to induce ovulation. After a period of 48-52 hours, these mice received an intraperitoneal injection of 5 IU human chorionic gonadotropin (hCG). Approximately 15-17 hours post hCG injection, the female mice were euthanized. Their fallopian tubes were aseptically extracted and placed in a culture dish containing a fertilization solution, which was then covered with mineral oil to minimize contamination. The ampullary region was punctured using a dissecting needle to release the oocytes into 0.2 mL of human tubal fluid (HTF) medium.

The male PWK/PhJ mice, originally purchased from the Jackson Laboratory (Bar Harbor, USA), were used to provide mature sperm cells. Sperms were collected from 12-week-old PWK/PhJ male mice by squeezing the cauda epididymis. The sperms were then placed in mineral oil containing FERTIUP Mouse Sperm Preincubation Medium and incubated at 37°C for 30 minutes. Subsequently, approximately 5 μL of the sperm suspension was added to a droplet of HTF medium containing the oocytes, resulting in a final concentration of approximately 200-600 sperms/mL. The culture dish was incubated at 37°C with 5% CO_2_ for 5-6 hours.

Following incubation, the dish was washed twice with fresh HTF medium. The embryos were then cultured in a CO_2_ incubator at 37°C and 5% CO_2_ for 18-22 hours. The embryos were washed with M2 medium and placed into droplets containing an appropriate number of embryos. Finally, the embryos were gently transferred into the expanded region of a surrogate C57BL/6J female mouse’s fallopian tube using a pipette. All animal maintenance and experimental procedures were conducted in strict accordance with the guidelines of the Institutional Animal Care and Use Committee (IACUC) of Tsinghua University, Beijing, China.

### Tissue digestion of F1 hybrid mice

F1 mice from a C57BL/6J × PWK/PhJ cross were maintained under controlled conditions (23±2°C, 55±15% humidity, 12/12 dark/light cycle) with ad libitum food and water. Animal experiments were performed in accordance with the ARRIVE (Animal Research: Reporting of In Vivo Experiments) guidelines and approved by Tsinghua University’s Institutional Animal Care and Use Committee (Approval No. 17-XW1). F1 mice were euthanized via cervical dislocation, sanitized with 75% alcohol, after which their lungs and livers were extracted and minced. The tissue fragments were then washed in ice-cold HBSS and chilled. Resulting cell suspensions were filtered (40µm Nylon mesh), centrifuged (300×g, 5 minutes), and the supernatant was discarded. Cells were then digested with 0.25 mg/mL Liberase TM (Roche, Basel, Switzerland) for 10 minutes. Following a double phosphate-buffered saline (PBS) wash, cells were resuspended in fresh culture medium as single-cell suspensions for subsequent experiments.

### Crosslinking and lysis of cells

Cell crosslinking, lysis and following DNA purification to sequencing steps were the same for human and mouse cells. Approximately 10 million cells were resuspended in 10 mL of fresh culture medium. Formaldehyde (278 μL of 37%) was added for fixation and cells were incubated 10 minutes at room temperature (RT), followed by the addition of 894 μL if 2.5 M glycine to halt the reaction. After a further 5 minutes RT incubation and 10 minutes on ice, cells were centrifuged (1000×g, 5 minutes, 4°C), washed twice with 5 mL of ice-cold 1× PBS, and the pellet was stored at −80°C.

For cell lysis, 3 million crosslinked cells were resuspended in 1 mL of ice-cold Hi-C lysis buffer (10 mM Tris-HCl pH 7.5, 10 mM NaCl, 0.2% NP-40, and 1× Roche protease inhibitors Cat#11697498001) and rotated at 4 °C for 30 minutes. After centrifugation (1000×g, 5 minutes, 4°C), the supernatant was discarded, and the nuclear pellet was washed with 500 µL of Hi-C lysis buffer. The pellet was then resuspended in 50 µL of 0.5% SDS and incubated at 62 °C for 10 minutes without shaking. After adding 145 µL of water and 50 µL of 10% Triton X-100, samples were rotated at 37 °C for 15 minutes to quench SDS. Subsequently, 25 µL of NEB (New England Biolabs, Ipswich, USA) Buffer 3.1 and 10 µL of 10 U/µL DpnII restriction enzyme (Cat#R0543T, NEB) were added, with a 4 hours rotation at 37 °C. After heat inactivation of DpnII (62 °C, 20 minutes), samples were rotated at 4 °C for 5 minutes. Finally, ligation master mix (750 µL) comprising of 100 µL of 10× NEB T4 DNA ligase buffer with 10 mM ATP, 75 µL of 10% Triton X-100, 3 µL of 50 mg/mL BSA (Cat#AM2616, Thermo Fisher Scientific, Waltham, USA), 10 µL of 400 U/µL T4 DNA Ligase (Cat#M0202, NEB), and 562 µL of water was added. Reactions were then rotated at 16 °C for 4 hours and at room temperature for 1 hour.

### DNA purification

For ordinary Pore-C library construction, the following protocol was followed: (1) Crosslink reversal: 45 µL of 10% SDS and 55 µL of 20 mg/ml Proteinase K were added and incubated at 63°C for a minimum of 4 hours, preferably overnight. (2) Addition of 65 µL of 5 M NaCl and incubation at 68°C for 2 hours. (3) Addition of 20 µL of 20 mg/mL RNase and incubation at 37°C for 1 hour. (4) Mixture of phenol:chloroform:isoamyl alcohol 25:24:1 (500 µL) was added and thoroughly mixed. The mixture was then transferred to a 2 mL MaXtract High Density tube (Cat#129056, Qiagen, Hilden, Germany) for phase separation. (5) A combination of 1 µL GlycoBlue (Cat#AM9515, Thermo Fisher Scientific), 100 μL of 3 M sodium acetate (pH 5.2), and 850 μL of isopropanol was added and the mixture was incubated at −80°C for 1 hour. (6) The mixture was centrifuged at maximum speed for 30 minutes at 4°C and the supernatant was discarded. (7) The pellet was washed twice with ice-cold 75% ethyl alcohol and the dried pellet was resuspended in 50 μL of Buffer EB.

For high throughput Pore-C library construction, DNA purification was performed the same as Pore-C except for an additional pronase digestion step. The dried pellet in the 7^th^ step above was resuspended in a solution containing 800 μL of ddH_2_O, 50 μL of 10% SDS, 100 μL of 1M Tris, and 50 μL of 20 mg/mL pronase (Cat#10165921001, Roche), followed by incubation at 40°C for 1 hour. Then 300 µL of phenol:chloroform:isoamyl alcohol (25:24:1) was added and mixed thoroughly. After centrifugation (maximum speed, 30 minutes, 4°C), the aqueous phase was collected, 90 μL of 3 M sodium acetate (pH 5.2) and 800 μL of isopropanol were added, and the mixture was incubated at −80°C for 1 hour. A subsequent centrifugation (maximum speed, 30 minutes, 4°C) was performed, supernatant was discarded, and the pellet was washed twice with ice-cold 75% ethyl alcohol before being resuspended in 50 μL of Buffer EB.

### Preparation and sequencing of Pore-C libraries

The ONT (Oxford Nanopore Technologies, Oxford, UK) sequencing library was prepared using 3-4 μg purified DNA per sample. DNA fragments over 3 kb were size selected using the PippinHT system (Sage Science, Beverly, USA). The NEB Next Ultra II End Repair/dA-tailing Kit (Cat#E7546, NEB) was utilized for end repair and dA addition, followed by adapter ligation using the SQK-LSK109 kit (ONT). The DNA library quantity was evaluated using Qubit 4.0 fluorometer (Invitrogen, Carlsbad, USA). Approximately 700 ng library DNA per flowcell were subjected to sequencing on the ONT PromethION platform by the Genome Center of Grandomics (Wuhan, China). We generated 8 flowcells of Pore-C data for HG001, 4 flowcells of Pore-C for HG002, and 2 flowcells of Pore-C data for mixed DNA of the F1 mice. Raw sequencing signals were transformed into DNA sequences using the model ‘dna_r9.4.1_450bps_hac_prom.cfg’ in Guppy v4.5.3 software (ONT). Reads with average base qualities <7 (corresponding to <80% accuracy) were filtered.

### Pore-C read decomposition and reference mapping

We employed Falign to decompose Pore-C concatemers into fragments and map these fragments to the reference genome using the default parameters, with their original order information in reads preserved (Figure 1). GRCh38 (NCBI accession: GCA_000001405.15) and GRCm39 (GCA_000001635.9) were used as reference for the human datasets and F1 mice, respectively.

### Training Pore-C SNV calling model

We trained the Pore-C model for Clair3 on the basis of the pretrained model labeled ‘r941_prom_hac_g360+g422’ using randomly sampled HG002 training data (Supplementary Table 1) of three different depths, 30×, 60×, and 90×. SNVs in high-confident regions from HG002 Genome in a Bottle (GIAB) v3.3.2 Benchmark datasets were used as truth variants in training. Both the pileup calling model and full-alignment model were retrained (refer to Supplementary Note 2) according to the detailed guidelines of Clair3 (https://github.com/HKU-BAL/Clair3). We used the default parameters in training with a learning rate of 5E-4 and a total of ten epochs. The Pore-C model was trained on Clair3 v0.1-r7, which also works for Clair3 v0.1-r10 invoked by Dip3D.

### SNV calling and evaluation

SNV calling was carried out using Clair3 v0.1-r10 with the Pore-C model for human datasets. In evaluation, GIAB v3.3.2 gold standard SNVs of HG001 and HG002 were used as truth variants for them, respectively. Detailed commands used in SNV calling and evaluation are described in Supplementary Note 3.

### SNV phasing and evaluation

In SNV phasing step, we first used the ‘extractHAIRS’ command from HapCUT2 v1.3.1^26^ and the genome-wide variants (VCF format) to convert the fragment alignment (BAM) file to the compact fragment file format, which contains local haplotype information of individual fragments. Then we wrote a script ‘make-pore-c-frag-pair’ that retrieves pairwise contact information from the BAM into a paired fragment file. Finally, the compact fragment file, paired fragment file and the VCF file were subjected to haplotype assembly by HapCUT2 that output a phased VCF file.

We evaluated the switch and hamming error rates of SNV phasing using the ‘compare’ command of WhatsHap v1.2^27^. For HG001, the benchmark phased VCF from GIAB v3.3.2 was used as ‘truth’ VCF. For the F1 mice dataset, we downloaded SNVs of the two parental strains (C57B/6J and PWK/PhJ) from the Mouse Genomes Project^34^. Then we screened SNVs with different homozygous genotypes in C57B/6J and PWK/PhJ using the ‘SNPsplit_genome_preparation’ command of SNPsplit v0.6.0, based on which we obtained a hypothesized F1 hybrid phased VCF heterozygous on all these SNVs as ‘truth’ VCF.

For Hi-C based phasing, we downloaded 531× HG001 Hi-C data aligned to GRCh38 from the study of Rao et al.^3^, which used the restriction enzyme MboI (isoschizomers of DpnII) and an in-situ Hi-C protocol, similar to our DpnII and in-situ Pore-C protocol. The SNVs of HG001 from GIAB v3.3.2 with GRCh38 as reference were applied in Hi-C based phasing. For F1 mice, we downloaded Hi-C data (NCBI accession: SRR5122741 and SRR5122742) of a similar C57B/6J × PWK/PhJ strain from the study of Du et al.^10^. Bowtie2 v2.5.1^35^ was used to map Hi-C reads, and HapCUT2 v1.3.1 was used for Hi-C based phasing. Phasing evaluation was performed the same as for the Pore-C data. Detailed commands are described in Supplementary Note 4.

### Fragment haplo-tagging

We first compared the h-trans contact rates after haplo-tagging using WhatsHap v1.2 with default settings and our own script with different SNV mismatch-free context lengths, mapping qualities (MAPQs), and percentages of alignment identity (PI). Pairwise contacts with both fragments haplo-tagged were extracted from the tagged bams and subjected to h-trans statistics. H-trans contact rate was calculated by dividing the number of h-trans contacts by the total count of h-trans and h-cis contacts. According to assessment results, the haplo-tagging module implemented in Dip3D applies three default thresholds, only using SNVs with 6 bp mis-match free contexts on fragments, MAPQ ≥ 5, and PI ≥ 85%. For Hi-C, we used HaploHiC v0.34 for fragment haplo-tagging. Detailed commands are available in Supplementary Note 5.

For Hi-C data of F1 mice, we used SNPsplit v0.3.2, Bowtie2 v2.5.1 and HiCUP v0.9.2^36^ for fragment haplo-tagging, as previously described^37^. Briefly, we bulit a N-masked GRCm39 genome using SNPsplit v0.3.2, where the SNVs sites of the two parental strains (C57B/6J and PWK/PhJ) were masked by the ‘N’ base. The Hi-C paired end reads were aligned to the N-masked reference using Bowtie2, and the alignments were further classified and filtered using HiCUP. Finally, the reads with overlapping SNVs sites were assigned to a certain allele using SNPsplit. Detailed commands are available in Supplementary Note 6.

### Haplotype imputation and final output

For haplotype imputation, we made multiple statistics to learn the characteristics of haplotype fragment distribution in Pore-C reads. We first studied the relationship between h-trans rate and the fragment interaction distance for all the contacts and the adjacent contacts respectively (Figure 5A). For the distance rule, interaction distances with 5% h-trans ratios were determined as the distance thresholds for distinguishing short-range and long-range interactions. To check the viability of the bridging rule, we calculated the ratio of *1*2*1* and *2*1*2* type reads among all reads with at least three haplo-tagged fragments. Here, 1 and 2 denote fragments from homologous chromosomes and * denotes any number (including zero) of fragments from any haplotype. For the dominant haplotype rule, we calculated the total ratio of minor-haplotype (occupying ≤1/2 haplo-tagged fragments in a h-trans read) fragments among all fragments in fully haplo-tagged reads containing ≥3 fragments.

After haplotype imputation, haplotype-unassigned fragments were dumped, haplotype-assigned fragments in h-cis reads were directly grouped into h-cis fragment groups, and h-trans reads were split into separate h-cis fragment groups (containing ≥2 h-cis fragments) based on haplotypes. All h-cis fragment groups are stored in a specialized SAM format that includes both the haplotype information and high order information of the fragments. Dip3D also provides several supplementary tools for extracting diploid contact matrix and haplotype-specific high order interaction analyses using the above output. For Hi-C, haplotype imputation was carried out using HaploHiC v0.34^13^. Detailed commands and algorithms for haplotype imputation are described in Supplementary Note 5.

### Assessments of diploid contact matrices

For evaluation, we compared Hi-C and Pore-C based diploid contact matrices. For Hi-C data of HG001, we downloaded the diploid contact matrix based on 2.6Tb (∼877.5×) Hi-C data from the study of Rao et al.^3^ that used GRCh37 as reference. For consistency, we applied Dip3D on 190× Pore-C data with GRCh37 as reference. We used Dip3D to extract h-cis pairwise contacts from the haplo-tagged fragment BAM for compatibility with Hi-C analytic tools. The command ‘pre’ of juicer-tools v1.22.01^38^ was used to convert the pairwise contacts into a diploid contact matrix in hic format. We used the command ‘hicConvertFormat’ in HiCExplorer v3.6^39^ to transform the hic matrix to the cool format. Cooler v0.8.11^40^ was used for matrix balancing. The diploid hic and cool files were used for diploid contact matrix visualization and calculation, respectively.

We used the same contact map resolution definition raised by Rao et al^3^: a diploid 3D genome is considered to reach resolution of K bp, if over 80% of its K bp bins have contact counts no less than 1,000. We used QuASAR-QC^29^ in 3DChromatin_ReplicateQC v1.0.1^30^ for quality evaluation of individual contact matrices.

The comparison between Pore-C and Hi-C contact matrices were based on the derivative values calculated by cooltools v0.5.1 (https://github.com/open2c/cooltools). We randomly downsampled an equal amount (160M per parental haplotype) of haplotype-assigned Pore-C pairwise contacts before converting them to a diploid contact matrix. Correlations between diploid matrices were analysed on two levels: (1) A/B compartment based on eigenvector scores calculated by cooltools eigs-cis; (2) TAD structure based on insulation scores calculated by cooltools insulation. We used HiCRep^41^ to evaluate the correlation between maternal and paternal contact matrices. Detailed commands used in section are described in Supplementary Notes 7,8.

### Haplotype-specific high order chromatin interaction analysis of gene regulatory regions

We screened h-cis fragment groups with no less than 5 fragments from the HG001 data for haplotype-specific high order interaction analyses. The gene promoter regions were obtained based on the GRCh38 GENCODE v29 gene annotation (ENCODE acc. ENCFF159KBI) as the 2kb regions upstream and downstream of the gene transcription start sites. We then used Bedtools v2.28^42^ to select Pore-C h-cis fragment groups overlapping with promoter regions for following analyses.

The single-chromatin distal to proximal contact ratio (SDPR) resembles the concept of distal to local contact ratio (DLR) previously used on Hi-C^43^, except that SDPR calculates the ratio within a single-chromatin (a h-cis fragment group) rather than the ratio (DLR) of cumulative Hi-C contacts overlapping with the investigated region. SDPR is calculated the ratio of the number of fragments with >3Mb (distal) interaction distances from the target fragment (overlapped with a target regions such as promoter) to those with ≤3Mb (proximal) interaction distances on the chromosomes in a h-cis fragment group.

For single-molecule multiway enhancer interaction analysis, we obtained the enhancer regions from the Encode cRES v3 functional element annotation library, which were extended 1kb upstream and 1kb downstream. We used Bedtools v2.28 to identify Pore-C fragments overlapping with enhancer elements and then calculated the number of overlapped enhancers in each h-cis fragment group. The promoter overlapping h-cis fragment groups were then classified as NE (no enhancer interaction), SE (single-enhancer interaction), and ME (multi-enhancer interaction) that interact with ≥2 enhancers, respectively.

The gene expression quantification data of HG001 (Acc number: ENCFF678BLG, ENCFF897XES, ENCFF791MED, ENCFF473KMX) were downloaded from the Encode database. We obtained the gene lists of allele-specific expression genes^13^, autosomal imprinted genes^44^, genes subjected to XCI, and escape genes including those variable on chrX^45^ from previously published studies.

## Data Availability

The in-house Pore-C data of HG001, HG002 and F1 mice have been deposited in the Genome Sequence Archive (GSA) at the BIG Data Center, Beijing Institute of Genomics, Chinese Academy of Sciences (BIG, http://gsa.big.ac.cn), under the Project Accession No. “PRJCA018069”. The GSA-Human and GSA Accession Nos. for the data are “HRA004983” and “CRA011676”, respectively.

The Pore-C model for Clair3 is available at http://www.bio8.cs.hku.hk/porec/clair3_porec_model.zip. The ONT data of HG001 and HG002 are available at https://s3-us-west-2.amazonaws.com/human-pangenomics/index.html?prefix=NHGRI_UCSC_panel. The GIAB v3.3.2 ‘truth’ VCFs of HG001 and HG002 were obtained from https://ftp-trace.ncbi.nlm.nih.gov/giab/ftp/release/NA12878_HG001 and https://ftp-trace.ncbi.nlm.nih.gov/giab/ftp/release/AshkenazimTrio. The region stratifications of GRCh38 reference genome are available at https://github.com/genome-in-a-bottle/genome-stratifications.

The processed Hi-C data of HG001 were obtained from Rao et al. (2014), including diploid Hi-C matrices (GSE63525) and 531× alignment files (https://data.4dnucleome.org/publications/cf0e49aa-173c-49d1-a7c7-22acbc83c064).

The SNV VCF files for the parental strains (C57B/6J and PWK/PhJ) of the F1 mice are available from the Mouse Genomes Project (https://ftp.ebi.ac.uk/pub/databases/mousegenomes/REL-2112-v8-SNPs_Indels/mgp_REL2021_snps.vcf.gz). Diploid HG001 Hi-C matrices from Rao et al.4 is available from GSE63525. The Hi-C data of C57B/6J × PWK/PhJ mice are available in NCBI SRA database under SRA accession IDs SRR5122741 and SRR5122742.

## Supplementary Figures

**Supplementary Figure 1.**
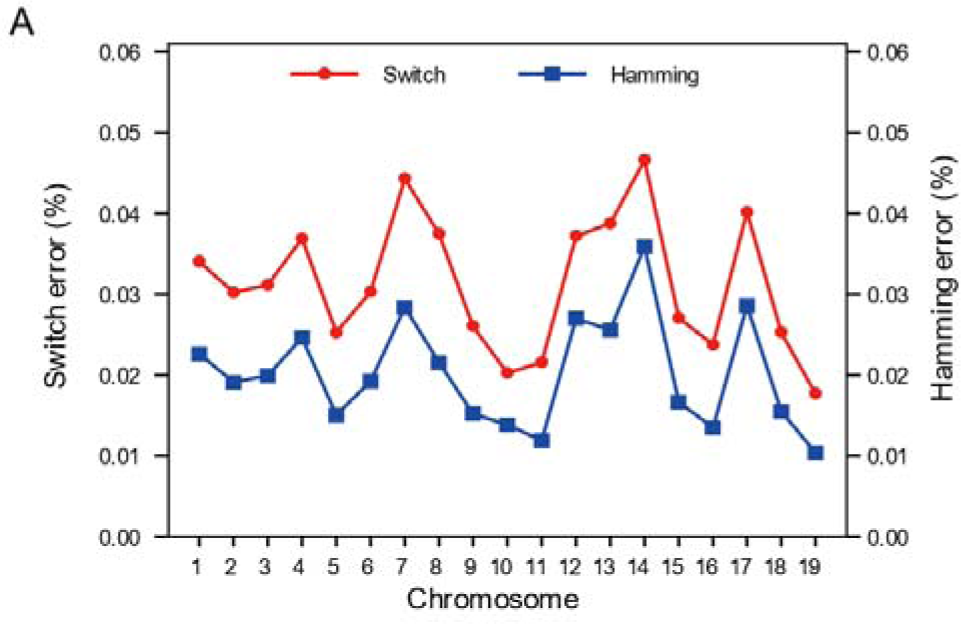
Switch and hamming error rates of Dip3D SNP phasing on the F1 mice Pore-C data.

**Supplementary Figure 2.**
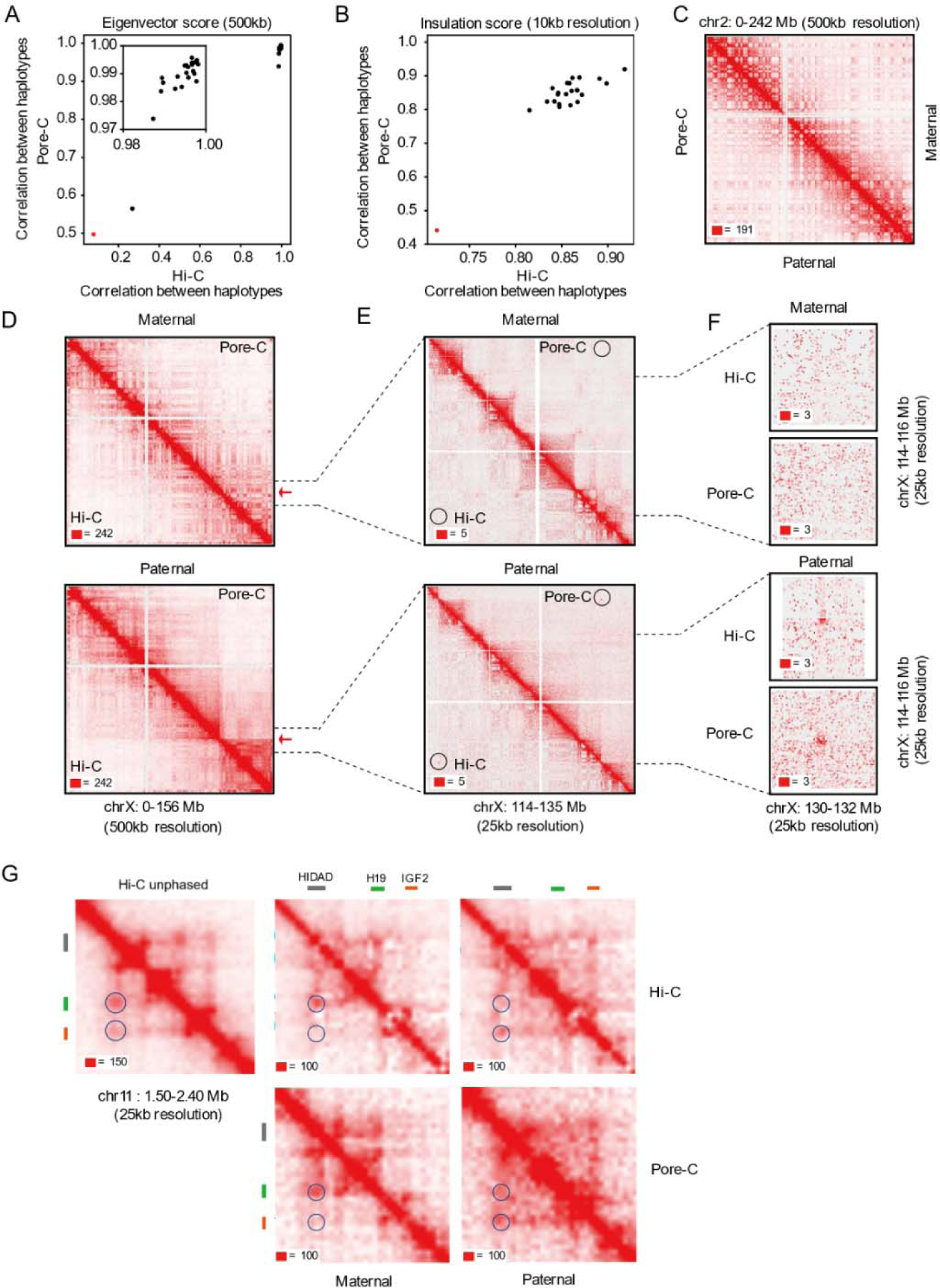
Reproduction of canonical diploid 3D genome structures in HG001 using Dip3D. **A** and **B**, corresponding correlation coefficients of A/B compartment eigenvector scores (A) and TAD insulation scores (B) between homologous chromosomes in Pore-C (Dip3D) and Hi-C diploid contact matrices. The black and red dots indicate values for autosomes and the X chromosome, respectively. **C**, Dip3D diploid contact matrices on an exemplary autosome (chr2). Red squares at the left bottom of the matrices indicate the color scale. **D**-**F**, Hi-C and Pore-C based diploid contact heatmaps of the entire chromosome X (D), the superdomain (at ∼115Mb) region (E), and the superloop (formed between tandem repeats DXZ4 and FIRRE) region (F). The superloop shown in F is also highlighted using circles in panel E. **G**, Hi-C and Pore-C diploid contact heatmap at the H19/Igf2 Distal Anchor Domain (HIDAD).

**Supplementary Figure 3.**
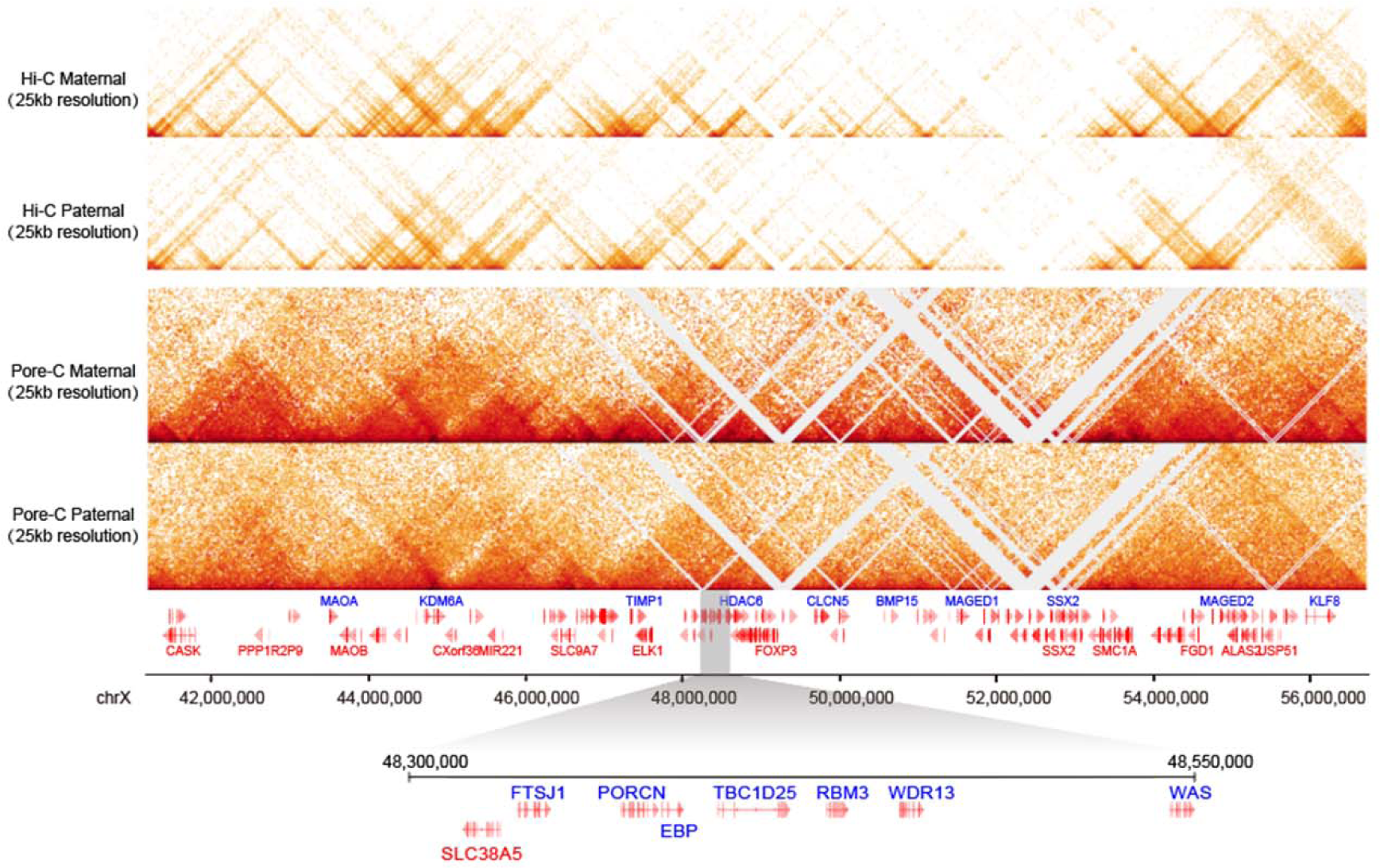
An exemplary Hi-C contact-sparse region with haplotype-specific structures visible in Dip3D matrices. Matrices based on 878× Hi-C data and 190×Pore-C data (Dip3D) are shown in this graph. The grey shadow indicates a previously reported region containing multiple allele-specific expression genes, which also shows significantly different haplotype-assigned contact frequencies between paternal and maternal haplotypes in the HG001 Dip3D matrices.

**Supplementary Figure 4.**
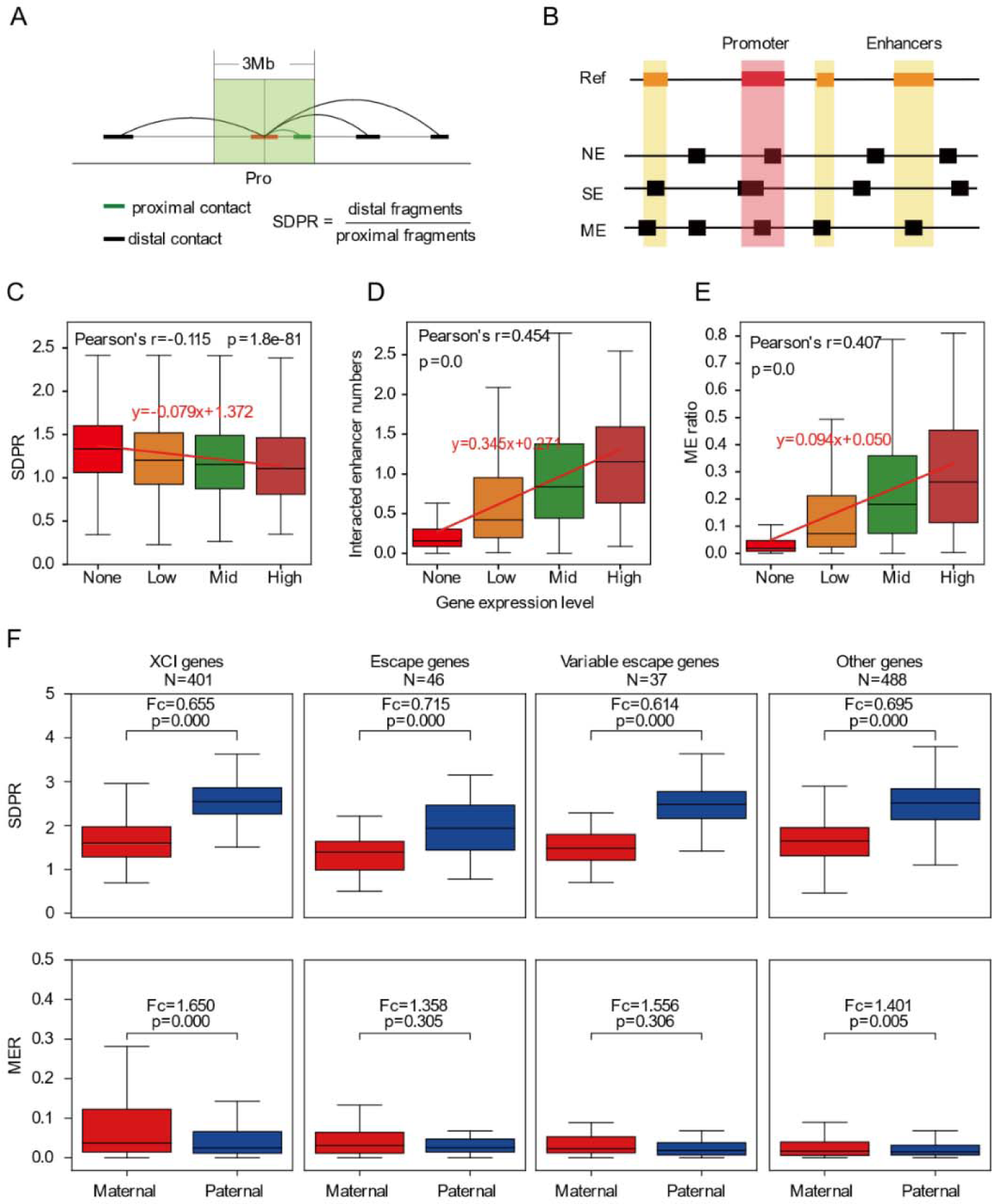
Statistics of promoter high order interactions and allelic differences. **A**, diagram showing the calculation of single-chromatin distal to proximal contact ratio (SDPR). **B**, different patterns of promoter-enhancer interaction observed in high-order reads. NE, SE, and ME denote the promoter interacts with zero, a single, and multiple enhancers in a high order read, respectively. **C**-**E**, correlations between gene expression levels (horizontal axes) and genic promoter SDPR (C), promoter-interacting enhancer number (D), and ME read ratio (E). Genes were ranked ascendingly according to their expression amount in HG001 transcriptomes and classified into four categories: None, zero expression; Low, ≤75%; Mid, >75% and ≤99%; High, >99% rank. **F**, distribution of allelic SDPRs (top) and allelic multi-enhancer interacting read ratios (MERs, bottom) among X chromosome genes subjected to X chromosome inactivation (XCI) and those escape from it. The classification of X chromosome genes was based on the study of Balaton et al., (2015). XCI genes and escape genes denote genes constantly subjected to XCI and XCI escape in different human cell lines, respectively. Variable escape genes only escape from XCI in a proportion of human cell lines. The status of the other genes had not been fully understood.

