## Supplementary Notes for "High-resolution diploid 3D genome reconstruction using Pore-C data"

#### Supplementary Note 1: Pore-C sequencing for HG001, HG002 and F1 mice

##### Human datasets

We generated 8 flowcells (6 ordinary and 2 high throughput) of Pore-C data for human cell line HG001 (GM12878) and 4 flowcells of (1 ordinary and 3 high throughput libraries) Pore-C data for HG002 (GM24385). The specifics for each flowcell, including read count, data yield, mean read length, N50 read length, mapped read count, fragment number and length, and pairwise contacts, are enumerated in Supplementary Table 1. The read counts of data passing the quality filter (average base quality) fluctuated between 4.5 million and 40.8 million, corresponding to data yields from 35.5 Gb to 143.6 Gb. Mean read lengths and N50 read lengths range from 3.0 kb to 7.8 kb and 4.9 kb to 9.8 kb, respectively. Moreover, the mean fragment lengths and the number of fragments per read ranged from 723.7 bp to 1127.8 bp and 3.3 /read to 8.5 /read, respectively. Pairwise contacts ranged from 139 million to 645 million. Among them, high throughput Pore-C flowcells have better sequence yield and more pairwise contacts than Pore-C flowcells.

##### F1 mice datasets

We sequenced 2 flowcells of Pore-C data using the high throughput library construction strategy for the F1 mice (C57BL/6J × PWK/PhJ) (see Methods). Detailed statistics for the two flowcells are available in Supplementary Table 1. The clean (quality-filtered) data read counts were 15.4 million and 19.2 million, and the corresponding data yields were 63.5 Gb and 93.7 Gb. Their mean read lengths reached 4.1 kb and 4.3 kb, with 4.3 kb and 4.8 kb read length N50. The mean fragment lengths were 811.2 bp and 817.0 bp, and on average they had 4.7 and 5.1 fragments per read, respectively. Pairwise contacts converted from the high-order reads were 186 million and 294 million.

##### Supplementary Node 2: Training Pore-C SNV model for Clair3

In this study, we trained the Pore-C SNV model for Clair3 (tested on v0.1-r7 and v0.1-r10) using data of human cell line HG002, including both the pileup and full-alignment models. We randomly subsampled 30× and 60× data from the total 90× HG002 data. Variants from the HG002 GIAB v3.3.2 benchmark dataset in the high-confidence regions (provided by GIAB bed file) were downloaded as candidate truth sets. We applied the Representation Unification algorithm to the 90× bam file to filter out candidate truth variants with low minor-allele frequency (<0.08) or inconsistent with the bam file. The filtered SNVs were used as the final true variants in model training and finetuning at different Pore-C sequencing depths.

To train the pileup model, we selected positive and negative samples from chr1-19, chr21, and chr22. The candidate minor allele frequency thresholds for SNV and small indel calling were set at 0.08 and 0.15 respectively. The non-variants were also selected within the high-confidence regions (bed file from GIAB). We disabled the setting parameter MAXIMUM_NON_VARIANT_RATIO by setting it to ‘None’ to use all screened truth variants and non-variants in pileup model training. We used Clair3 default training parameters for the rest settings.

To train the full-alignment model, we selected all variant calls from calling results of the pileup model using the three BAMs with 30×, 60×, and 90× depths, as well as the top 30% reference calls sorted by quality scores for fine-tuning. We set the MAXIMUM_NON_VARIANT_RATIO to 10, limiting the maximum ratio of used variants to non-variants in training to 1:10. We used Clair3 default training settings for the other parameters.

We trained the Pileup and Full-alignment models based on the Clair3 r941_prom_hac_g360+g422 WGS models at a learning rate of 5e-4 with a total 10 epochs. To test the model's performance, we used 90× randomly sampled HG001 Pore-C dataset. Before calling SNVs, we first filtered out alignments with alignment length (inferred from SAM cigar string) below 100 bp, which are significantly enriched in poor quality alignments than the longer alignments. After filtering, ~81× of test data were left. Variant calling was performed using Clair3 (version v0.1-r8) with the newly trained models. We used hap.py (v0.3.12) to validate the calling result with the GIAB HG001 v3.3.2 truth variants. Testing results on HG001 and HG002 data are listed in Table 1.

#### Supplementary Node 3: Pore-C SNV calling and evaluation

Dip3D calls SNVs for Pore-C data with Clair3 (v0.1-r10):

docker run -it \

-v "${snp_ref_dir}":"/ref_dir" \

-v "${clair3_output}":"/output_dir" \

-v "${snp_bam_dir}":"/input_dir" \

-v "${snp_model_dir}":"/model_dir" \

hkubal/clair3:"${CLAIR3_VERSION}" \

/opt/bin/run_clair3.sh \

--bam_fn=/input_dir/snp-frag-ref.bam \

--ref_fn=/ref_dir/reference.fa \

--threads=${I_threads} \

--platform="ont" \

--model_path=/model_dir \

--output=/output_dir \

--var_pct_full=0.1 \

--call_snp_only

The variants output by Clair3 is

dip3d-output/2-snp/merge_output.vcf.gz

To evaluate SNV calling with HG001 Pore-C dataset, we used the GIAB v3.3.2 calling set downloaded from

<https://ftp-trace.ncbi.nlm.nih.gov/giab/ftp/release/NA12878_HG001/NISTv3.3.2/GRCh38/HG001_GRCh38_GIAB_highconf_CG-IllFB-IllGATKHC-Ion-10X-SOLID_CHROM1-X_v.3.3.2_highconf_PGandRTGphasetransfer.vcf.gz>

as truth SNVs.

To evaluate SNV calling with HG002 Pore-C dataset, we use the GIAB v3.3.2 calling set downloaded from

<https://ftp-trace.ncbi.nlm.nih.gov/giab/ftp/release/AshkenazimTrio/HG002_NA24385_son/NISTv3.3.2/GRCh38/HG002_GRCh38_GIAB_highconf_CG-Illfb-IllsentieonHC-Ion-10XsentieonHC-SOLIDgatkHC_CHROM1-22_v.3.3.2_highconf_triophased.vcf.gz>

as ground truth SNVs.

Both ground truth datasets are called for the GRCh38 reference genome available at

<ftp://ftp.ncbi.nlm.nih.gov/genomes/all/GCA_000001405.15_GRCh38/seqs_for_alignment_pipelines.ucsc_ids/GCA_000001405.15_GRCh38_no_alt_plus_hs38d1_analysis_set.fna.gz>

As described in Clair3, we generated high-confidence regions for the two truth datasets by removing the GA4TH low-complexity regions

AllRepeats_lt51bp_gt95identity_merged.bed

AllRepeats_51to200bp_gt95identity_merged.bed

AllRepeats_gt200bp_gt95identity_merged.bed

SimpleRepeat_imperfecthomopolgt10_slop5.bed

remapped_superdupsmerged_all_sort.bed

remapped_PacBio_MetaSV_svclassify_mergedSVs.bed

hg38_self_chain_nosamepos_withalts_gt10k.bed

which are available at

<https://github.com/jzook/genome-data-integration/tree/master/NISTv3.3.2/filtbeds/GRCh38>

GIAB’s high-confidence regions for HG001 available at

<https://ftp-trace.ncbi.nlm.nih.gov/giab/ftp/release/NA12878_HG001/NISTv3.3.2/GRCh38/HG001_GRCh38_GIAB_highconf_CG-IllFB-IllGATKHC-Ion-10X-SOLID_CHROM1-X_v.3.3.2_highconf_nosomaticdel_noCENorHET7.bed>

and for HG002 available at

<https://ftp-trace.ncbi.nlm.nih.gov/giab/ftp/release/AshkenazimTrio/HG002_NA24385_son/NISTv3.3.2/GRCh38/HG002_GRCh38_GIAB_highconf_CG-Illfb-IllsentieonHC-Ion-10XsentieonHC-SOLIDgatkHC_CHROM1-22_v.3.3.2_highconf_noinconsistent.bed>

The resulting high-confidence regions hg001_grch38.clean.bed for HG001 are generated via

VCF_HIGH_CONF_BED="/data1/cy/data/rna/grch38/hg001_grch38/HG001_GRCh38_GIAB_highconf_CG-IllFB-IllGATKHC-Ion-10X-SOLID_CHROM1-X_v.3.3.2_highconf_nosomaticdel_noCENorHET7.bed"

cat ${VCF_HIGH_CONF_BED} | \

bedtools subtract -a stdin -b AllRepeats_lt51bp_gt95identity_merged.bed | \

bedtools subtract -a stdin -b AllRepeats_51to200bp_gt95identity_merged.bed | \

bedtools subtract -a stdin -b AllRepeats_gt200bp_gt95identity_merged.bed | \

bedtools subtract -a stdin -b SimpleRepeat_imperfecthomopolgt10_slop5.bed | \

bedtools subtract -a stdin -b remapped_superdupsmerged_all_sort.bed | \

bedtools subtract -a stdin -b remapped_PacBio_MetaSV_svclassify_mergedSVs.bed | \

bedtools subtract -a stdin -b hg38_self_chain_nosamepos_withalts_gt10k.bed | \

bedtools subtract -a stdin -b GCA_000001405.15_GRCh38_no_alt_plus_hs38d1_analysis_set_REF_N.bed > hg001_grch38.clean.bed

The high-confidence regions hg002_grch38.clean.bed is generated in the same way. The evaluations are then performed on these two bed files with hap.py (v0.3.12):

REF_DIR="/data1/cy/data/rna/grch38/reference/"

INPUT_DIR="/data2/cy/data/snp/snp-output"

GT_DIR="/data1/cy/data/rna/grch38/hg001_grch38/"

GT_VCF="HG001_GRCh38_GIAB_highconf_CG-IllFB-IllGATKHC-Ion-10X-SOLID_CHROM1-X_v.3.3.2_highconf_PGandRTGphasetransfer.vcf.gz"

BED="hg001_grch38.clean.bed"

OUTPUT_DIR="/data2/cy/data/snp/hap-out3"

NT=8

docker run \

-v "${REF_DIR}":"/ref_dir" \

-v "${GT_DIR}":"/gt_dir" \

-v "${INPUT_DIR}":"/input_dir" \

-v "${OUTPUT_DIR}":"/output_dir" \

jmcdani20/hap.py:v0.3.12 /opt/hap.py/bin/hap.py \

/gt_dir/${GT_VCF} \

/input_dir/merge_output.vcf.gz \

-f /gt_dir/$^1^ \

-r /ref_dir/GCA_000001405.15_GRCh38_no_alt_plus_hs38d1_analysis_set.fna \

-o /output_dir/happy \

--engine=vcfeval \

--threads=${NT} \

--pass-only

The benchmark results with difference coverage of Pore-C and Hi-C data are listed in Table 1 and Supplementary Table 2.

To stat SNPs called on difficult regions, we download difficult regions including GC content, simple repeats, tandem repeats, segmental duplications and low mappability, from <https://ftp-trace.ncbi.nlm.nih.gov/giab/ftp/release/genome-stratifications/v2.0/GRCh38/>. These regions are provided in BED format and stored in directory /data2/linzhuobin/Templates/evaluation/genome-stratifications. We finally run the command

REF_DIR="/data1/chenying/dip3d-1/"

GT_DIR="/data2/linzhuobin/Templates/evaluation/giab"

VCF_DIR="/data2/linzhuobin/Templates/dip3d_COV/90X/results/2-snp"

BED_DIR="/data2/linzhuobin/Templates/evaluation/giab"

STRATIFICATION_DIR="/data2/linzhuobin/Templates/evaluation/genome-stratifications"

OUTPUT_DIR="${VCF_DIR}/happy"

docker run \

-v "${REF_DIR}":"/ref_dir" \

-v "${GT_DIR}":"/gt_dir" \

-v "${VCF_DIR}":"/vcf_dir" \

-v "${BED_DIR}":"/bed_dir" \

-v "${STRATIFICATION_DIR}":"/stratification_dir" \

-v "${OUTPUT_DIR}":"/output_dir" \

jmcdani20/hap.py:v0.3.12 /opt/hap.py/bin/hap.py \

/gt_dir/HG001_GRCh38_GIAB_highconf_CG-IllFB-IllGATKHC-Ion-10X-SOLID_CHROM1-X_v.3.3.2_highconf.vcf.gz \

/vcf_dir/all.phased_snp.vcf.gz \

-f /bed_dir/hg001_grch38.clean.bed \

-r /ref_dir/GCA_000001405.15_GRCh38_no_alt_plus_hs38d1_analysis_set.major.fna \

-o /output_dir/stratification \

--stratification /stratification_dir/v2.0-GRCh38-stratifications.tsv \

--pass-only \

-threads=20

The statistical results are given in Supplementary Table 2. Overall, 90X Pore-C generates 61.09% and 51.2% SNPs on these regions than 90X and 531X Hi-C respectively.

### Supplementary Note 4: Pore-C SNV phasing evaluations

Dip3D outputs phased SNV set in the path

dip3d-output/2-snp/phased_snp.vcf.gz

We use the following command to examine the switch error rate and Hamming error rate:

SNP_GT=/data1/chenying/dip3d/giab_hg001_grch38/HG001_GRCh38_GIAB_highconf_CG-IllFB-IllGATKHC-Ion-10X-SOLID_CHROM1-X_v.3.3.2_highconf_PGandRTGphasetransfer.vcf

SNP_CALL=/data1/chenying/dip3d/dip3d-output/2-snp/phased_snp.vcf.gz

whatshap compare --ignore-sample-name --tsv-pairwise compare.pw.tsv --longest-block-tsv compare.block.tsv --only-snvs ${SNP_CALL} ${SNP_GT}

We use the following command to examine phasing completeness and resolution:

whatshap stats dip3d-output/2-snp/phased_snp.vcf

The phasing evaluation results are listed in Supplementary Table 3. With 30x Pore-C achieved an average whole-genome phased SNV density of 0.72/kb. Further increment of Pore-C data depth to 60x and 90x only resulted in 1.65-2.18% improvement in SNV phasing rate and increased the phased SNV density by 1.06-1.08 times respectively, suggesting that a 30x depth of Pore-C is near saturation.

We next consider phasing completeness, that is, the spanning of most-variants-phased (MVP) blocks on each chromosome. From Supplementary Table 3 we can see that Pore-C generates chromosome-level MVP phase blocks on all 23 chromosomes with only 30× data. The 23 chromosomes achieve phasing completeness (The phasing completeness of a chromosome is the proportion of bases spanned by its MVP block) as high as 79.3-100%. A deeper analysis shows that the low phasing completeness is caused by the widely existence of gap regions (a gap region, or a gap for short, is an ambiguous subsequences filled by N’s) in the reference genome. For example, as high as 27.1% of chr22 is occupied by gaps. Since these large gaps are not covered by any Pore-C reads, the MVP block of chr22 only covers 79.3% of the bases. Nevertheless, the MVP block still spans the whole non-gap regions of chr22, meaning that all of the non-gap regions of chr22 are in phase.

We next consider phasing resolution (the phasing resolution of a chromosome is defined as the proportion of phased SNVs contained in its MVP block). The resolutions of the 23 chromosomes are 97.19-98.94% with 30X Pore-C and rise to 98.49-99.60% with 90X Pore-C.

We further assessed the accuracy of the phasing results by scrutinizing switch and hamming errors. From Supplementary Table 3, it is seen that the accuracy increases as we improve the data depth. The switch error and hamming error of 30x Pore-C are 0.59% and 0.44% respectively and decrease to 0.36 and 0.28 respectively with 90x Pore-C.

To compare the phasing performance between Hi-C and Pore-C, we conduct a re-phasing of the GIAB HG001 variant sets (V3.3.2). We first extract the heterozygous variants with the following commands:

bcftools view -v snps,indels -m 2 -M 2 -O z -o HG001.snvs.vcf.gz HG001_GRCh38_GIAB_highconf_CG-IllFB-IllGATKHC-Ion-10X-SOLID_CHROM1-X_v.3.3.2_all.vcf.gz

bcftools annotate -x INFO,^FORMAT/GT -O z -o HG001.snvs.GT.vcf.gz HG001.snvs.vcf.gz

bcftools filter -i ' FILTER=="PASS" ' -O z -o HG001.snvs.GT.PASS.vcf.gz HG001.snvs.GT.vcf.gz

We next remove the genotype information:

gunzip -c HG001.snvs.GT.PASS.vcf.gz > HG001.snvs.GT.PASS.vcf

whatshap unphase HG001.snvs.GT.PASS.vcf > HG001.snvs.GT.PASS.unphased.vcf

We use Pore-C data to phase this unphased variants set in the same way as using Dip3D called SNVs. We also use Hi-C data to phase this set with in standard way. The results are given in Supplementary Table 3. For both Hi-C and Pore-C, the phasing completeness and phasing accuracy is improved as we continuously increase the data depth.

Phasing results using GIAB SNV sets were similar to that using the Dip3D called SNVs. The performance of Hi-C relies heavily on the data coverage. From Supplementary Table 3 we can see that although both Hi-C and Pore-C generates MVP blocks with the same spanning, Pore-C phases much more variants than Hi-C. For example, with 30x coverage, Pore-C phases 96.16-97.37% variants, while Hi-C only phases 51.5-70.0% variants. Even with 531x, Hi-C only phases 81.3-84.4% variants, which is significantly lower than 30x of Pore-C.

The switch errors and hamming error rates of Pore-C using the GIAB SNV sets is slightly higher than using Dip3D called SNVs (Supplementary Table 3). 30x Hi-C introduces much high hamming error rates because only 51.5-70% variants are phased. For example, on Chromosomes 1 and 2, the hamming error rates are as high as 48.9% and 41.05% respectively. Pore-C’s performance on phasing accuracy is better than Hi-C in coverage 30x, 60x and 90x. The phasing with 531x Hi-C achieves the best phasing accuracy.

The above evaluations show that using 90X Pore-C, the SNVs are phased with high accuracy. All the non-gap reference regions are in phase. And over 98.4% of the phased SNVs can be used for Pore-C fragment haplo-tagging.

### Supplementary Note 5: Pore-C fragment haplo-tagging

In Hi-C data, a mate-rescue strategy is applied for inferring the unknown haplotype for Hi-C cis interactions (between two Hi-C fragments on the same chromosome). Since the probability of h-trans error (interactions between two Hi-C fragments from homologous chromosomes rather than the same chromosome) $P(x)$ grows as the genomic distance of the interactions $x$ increase, the mate-rescue is only performed on interactions whose genomic distances are less than a threshold $L$. $L$ is usually set as the maximum value of $x$ such that $P(x)\leq0.05$. That is, $L:=max\{x:P(x)\leq0.05\}$. For example, for the mice Hi-C data, $L$ is set to 30 Mb and for human cell line HG001, $L$ is set to 50 Mb. In Dip3D, this idea, together with the ligation bridge characteristics of higher-order interactions, is also used for haplotype imputations. Specifically, Dip3D resolves haplotypes of the Pore-C fragments in three steps.

In the first step, the fragments are haplo-tagged based on their overlapped phased SNVs that generated in the previous step of Dip3D. We only used phased SNVs contained in the most-variants-phased (MVP) block of each chromosome to avoid introducing potential h-trans interactions between different phase blocks. We found that a directly fragment tagging with WhatsHap will result in h-trans error $P(x)$ greater than 0.05 on all genomic distances due to the high sequence errors. A deeper analysis reveals that $P(x)$, and the proportion of tagged fragments, depend on the number of mismatch-free bases surrounding the overlapped SNVs, the identity and the mapping quality of the fragments’ mapping results, as demonstrated in Figure 2. Based on these observations, we utilize two filtering strategies to achieve the best tradeoff between $P(x)$ and the number of tagged fragments. First, we only use overlapped SNVs that surrounded by 3 upstream match bases and 3 downstream match bases to infer the haplotypes. Second, those fragments whose mapping qualities are less than 5 or whose alignment identities are less than 85% will not be tagged in this stage. After this stage, the Pore-C reads are classified as completely phased, partially phased, and unphased reads.

In the second step, Dip3D resolves the value of threshold $L$. Based on the higher-order interactions captured in Pore-C reads, Dip3D employs two thresholds $L_{a}$ and $L_{g}$. $L_{a}$ is used for adjacent interactions and $L_{g}$ is used for other interactions. An interaction is called adjacent, if there exists no fragment that lie between the two fragments connected by this interaction in the read. On the human GM12878 Pore-C dataset, these two values are resolved to be 18 Mb and 20 Mb. They are also treated as the default threshold values in Dip3D for haplotype imputation.

In the final step three, a haplotype imputation is performed on partially phased reads. Suppose that after step one the fragments of a Pore-C read R=$f_{1}f_{2}\ldots f_{n}$ are partitioned into $x$ paternal fragments $P_{1},\ldots,P_{x}$, $y$ maternal fragments $M_{1},\ldots,M_{y}$ and $z$ unphased fragments $U_{1},\ldots{,U}_{z}$ such that $x+y+z=n$. Two fragments $f_{i}$ and $f_{j}$ are said to be close if either 1) $\left| i-j \right|=1$ and their genomic distance is less than $L_{a}$; or 2) $|i-j|\neq1$ and their genomic distance is less than $L_{g}$. For each paternal fragment $P_{i}$, we construct its rescue set ${CP}_{i}$ in the following way:

${CP}_{i}:=\{f_{k}: f_{k} \text{and} P_{i} \text{are close}\}$.

Based on the ligation bridging of intra-chromosomal fragments with spatial proximity, for each $f_{k}\notin{CP}_{i}$, we also add it to ${CP}_{i}$ if either $f_{k-1}$ or $f_{k+1}$ also belongs to ${CP}_{i}$. In this way we obtain $x$ paternal rescue sets ${CP}_{1},\ldots,{CP}_{x}$. We take the union of these $x$ paternal rescue sets into one paternal rescue set $CP$. In the same way we obtain a maternal rescue set $CM$. The fragments belonging to both $CP$ and $CM$ will generates h-trans interactions. We have to remove them from $CP$ and $CM$: $CP:=CP\backslash CM$, $CM:=CM\backslash CP$. Each unphased fragment in $CP$ is then imputed as paternal and each unphased fragment in $CM$ is imputed as maternal.

After the above imputation, if over 80% of this read’s fragments are tagged as paternal (maternal), then the remaining unphased fragments are all tagged as paternal (maternal).

Dip3D tags fragments on different chromosomes separately. Here we take chr1 as a running example. The tagging results with overlapped phased SNVs are given in

dip3d-output/3-frag-hap/chr1/snp-frag-hap.list

The tagging results after haplotype imputations are given in

dip3d-output/3-frag-hap/chr1/imputed-frag-hap.list

These two files have the same formats as shown below:

69	7	0	66504370	30	2

69	8	0	66496195	30	2

70	1	0	198379651	30	0

70	2	0	197737248	30	0

72	0	0	78145376	60	0

74	0	0	39638384	30	1

Each line corresponds to one fragment’s tagging results and has six columns. The meanings of the columns are: numeric read id, numeric fragment id, numeric chromosome id, the mapping position of this fragment, the mapping of quality of this fragment, and the haplotype of this fragment (0 for unknown, 1 for paternal and 2 for maternal). Furthermore, as is done in whatshap, Dip3D also outputs a BAM file containing haplotype information (HP:i:1 for paternal and HP:i:2 for maternal):

dip3d-output/3-frag-hap/chr1/tagged-chr1.bam

To analysis the tagging results, we extract corresponding information from these files. We first add mapping quality, mapping identity and fragment lengths to the tagging file:

samtools view ${chr}.bam |awk -F'\t|:|_' '{print int($2),int($3),int($4),int($5),$9,$21,length($14)}' |sort -n |tr ' ' '\t' > frag-map.list

We also need to add the surrounding bases at each overlapped SNVs:

zcat ${chr}.phased_snp.vcf |grep -v '^#'|awk '{print $1,$2-1,$2}'|tr ' ' '\t'|uniq > snp.bed

java -jar /home/zyserver/chenying/dip3d/dip3d/software/jvarkit/dist/sam2tsv.jar \

-R $REF_GENOME --regions snp.bed -N $BAM |\

awk '{if($10=="M") print $4,$8-1,$8,$1,$9,$6}'|tr ' ' '\t'|\

grep -C 5 -f snp.bed|grep -v '^--'|awk -v OFS='\t' '{print $4,$1,$3,$5,$6}' > adj5base.txt

zcat ${chr}.phased_snp.vcf|grep -v '^#'|awk -F'\t|:' '{if($14=="0|1") print $1,$2-1,$2,$4,$5;else print $1,$2-1,$2,$5,$4}'|tr ' ' '\t'|uniq > snp.bed

python /data3/linzhuobin/script/get_frag_snp.py snp.bed adj5base.txt frag-snp.list

For comparison, we also tag the fragments with whatshap:

samtools faidx GCA_000001405.15_GRCh38_no_alt_plus_hs38d1_analysis_set.fna

samtools index ${chr}.bam

whatshap haplotag ${chr}.phased_snp.vcf ${chr}.bam --ignore-read-groups\

-o Raw/tagged-chr1.bam --reference \

GCA_000001405.15_GRCh38_no_alt_plus_hs38d1_analysis_set.fna

samtools view Raw/tagged-chr1.bam |awk -F'\t|:|_' '{print int($2),int($3),int($4),int($5),int($24)}' |awk '!$5{$5="0"}1'|sort -n|tr ' ' '\t' > Raw/frag-hap.list

The tagging results output by whatshap (Raw/frag-hap.list), mapping information of fragment (frag-map.list) and the surrounding bases at overlapped bases are input to the script haplotag_characteristic_plot.ipynb to perform various analysis, such as h-trans error rates. We infer the distance thresholds $L_{a}$ and $L_{g}$, examine the effects of imputations with script dip3d_rescue.ipynb.

Supplementary Table 4 lists fragment and contact haplo-taggning statistics after each tagging and imputation step for Both Hi-C and Pore-C.

### Supplementary Note 6: Analysis of hybrid mouse data

We generated in house 60× Pore-C data for a F1 hybrid mouse (maternal C57B/6J and paternal PWK/PhJ). The SNVs are called and phased for the GRCm39 genome downloaded from

<https://ftp.ensembl.org/pub/release-109/fasta/mus_musculus/dna/Mus_musculus.GRCm39.dna.toplevel.fa.gz>

Decompress this file and build an index:

gunzip -c Mus_musculus.GRCm39.dna.toplevel.fa.gz > Mus_musculus.GRCm39.dna.toplevel.fa

samtools faidx Mus_musculus.GRCm39.dna.toplevel.fa

To evaluate the SNV calling and phasing performance of Pore-C, we download SNP sets corresponding to the two strains (C57B/6J and PWK/PhJ) from the Mouse Genomes Project:

<https://ftp.ebi.ac.uk/pub/databases/mousegenomes/REL-2112-v8-SNPs_Indels/mgp_REL2021_snps.vcf.gz>

We extract homozygous loci from this VCF file with SNPsplit (v0.6.0):

SNPsplit-0.6.0/SNPsplit_genome_preparation --vcf_file mgp_REL2021_snps.vcf.gz \

--strain PWK_PhJ --reference_genome GRCm39/ --genome_build GRCm39

With this command we obtain

all_SNPs_PWK_PhJ_GRCm39.txt.gz

which is further transformed to a standard phased VCF format：

gunzip -c all_SNPs_PWK_PhJ_GRCm39.txt.gz > all_SNPs_PWK_PhJ_GRCm39.txt

awk '{print $2,$3}' all_SNPs_PWK_PhJ_GRCm39.txt |sort -n -k1 -k2 |tr ' ' '\t'> phased.SNP.txt

bcftools view -s PWK_PhJ -T phased.SNP.txt mgp_REL2021_snps.vcf.gz -O v -o phased.SNP.vcf

python get_phased_vcf.py phased.SNP.vcf GT.phased.SNP.vcf

bgzip -c GT.phased.SNP.vcf > GT.phased.SNP.vcf.gz

tabix -p vcf GT.phased.SNP.vcf.gz

We run Dip3D on the 60× mouse Pore-C dataset, we evaluate the phasing performance:

\## Unify chrom_ID and sample_ID

cd /data3/linzhuobin/Templates/hybrid/compare

echo "C57M_PWKP" > sample.txt

sed 's/chr//g' phased_snp.vcf |bcftools reheader -s sample.txt -O z -o modchr.phased_snp.vcf.gz

tabix -p vcf modchr.phased_snp.vcf.gz

sed 's/chr//g' all.phased_snp.vcf | bcftools reheader -s sample.txt -O z -o modchr.all.phased_snp.vcf.gz

tabix -p vcf modchr.all.phased_snp.vcf.gz

\## Compare against benchmark SNPs

mkdir all mvp

whatshap compare --tsv-pairwise mvp/compare.tsv ../SNPsplit/GT.phased.SNP.vcf.gz modchr.phased_snp.vcf.gz > mvp/compare.log

whatshap stats --tsv mvp/stats.tsv modchr.phased_snp.vcf.gz > mvp/stats.log

To compare the interaction characteristics between Pore-C and Hi-C, we also downloaded a Hi-C dataset of another C57B/6J × PWK/PhJ strain (GSE82185, <https://www.nature.com/articles/nature23263>) from

<https://www.ncbi.nlm.nih.gov/geo/query/acc.cgi?acc=GSE82185>

We next transformed the data to FASTQ format:

fastq-dump --split-files SRR5122741 SRR5122742

gzip *.fastq

\## QC: remove adapter

trim_galore --paired --quality 20 --fastqc --illumina --length 20 -j 20 -o QC SRR5122741_1.fastq.gz SRR5122741_2.fastq.gz SRR5122742_1.fastq.gz SRR5122742_2.fastq.gz

ls QC

cat QC/*_val_1.fq.gz > HiC_hybrid_1.fastq.gz

cat QC/*_val_2.fq.gz > HiC_hybrid_2.fastq.gz

We preprocessed the reference genome and built and index for reference mapping with Bowtie2:

cut -f1 Mus_musculus.GRCm39.dna.toplevel.fa.fai > sort_chr.list

for chr in `cat sort_chr.list`;do cat PWK_PhJ_N-masked/chr${chr}.*fa >> GRCm39.N-masked.fa;done

bowtie2-2.5.1-linux-x86_64/bowtie2-build --threads 40 --large-index GRCm39.N-masked.fa GRCm39

We extracted the restriction enzyme sites from the reference genome:

HiCUP-0.9.2/hicup_digester --genome GRCm39 --re1 ^GATC,MboI GRCm39.N-masked.fa

The restriction enzyme sites are stored in

Digest_GRCm39_MboI_None_23-10-06_16-06-2023.txt

We edit the config file hicup_example.conf to fill in the paths of Hi-C reads, reference genome, restriction enzyme sites and run

HiCUP-0.9.2/hicup --config hicup_example.conf

To perform reference mapping and filtering of Hi-C. After mapping, we tag the BAM file with heterozygous SNPs generated above：

SNPsplit-0.6.0/SNPsplit --no_sort --hic --paired \

--snp_file all_SNPs_PWK_PhJ_GRCm39.txt HiC_hybrid_1_2.hicup.bam

We extract the tagging information of Hi-C reads for subsequent analysis：

samtools view -@ 40 HiC_hybrid_1_2.hicup.allele_flagged.bam |\

awk '{print $1,$3,$4,substr($NF,6,7)}'|tr ' ' '\t' > hic-hap.list

We feed the Pore-C tagging results (imputed-frag-hap.list) and Hi-C tagging results (hic-hap.list) into the script mouse_characteristic_plot.ipynb to analyze the interactions from different perspectives.

### Supplementary Note 7: Pore-C diploid contact maps evaluation

Dip3D tags Pore-C fragments on different chromosomes separately. Here we take chr1 as a running example. We first extract tagged records from the tagged BAM file:

samtools view -@40 -h -d HP:1 tagged-chr1.bam > HP1.sam

samtools view -@40 -h -d HP:2 tagged-chr1.bam > HP2.sam

and transform SAM format to PAF format:

paftools.js sam2paf HP1.sam > HP1.paf && rm HP1.sam

paftools.js sam2paf HP2.sam > HP2.paf && rm HP2.sam

Remove the suffices at read names to recover the original read names of each Pore-C fragment:

awk '{split($1,a,"_");print a[1],$2,$3,$4,$5,$6,$7,$8,$9,$10,$11,$12}' HP1.paf |sort -k 1,1|tr ' ' '\t' > tmp && mv tmp HP1.paf

awk '{split($1,a,"_");print a[1],$2,$3,$4,$5,$6,$7,$8,$9,$10,$11,$12}' HP2.paf |sort -k 1,1|tr ' ' '\t' > tmp && mv tmp HP2.paf

Transform PAF format to the standard jmatrix format of juicer tools:

python Generate_Contact_juiceMatrix_map_paf.py HP1.paf HP1.jmatrix

python Generate_Contact_juiceMatrix_map_paf.py HP2.paf HP2.jmatrix

Transform the jmatrix format to Hi-C cool format that can be visualization:

java -Xmx120g -jar juicer_tools_1.22.01.jar pre --threads 40 -r 5000,10000,25000,50000,100000,250000,500000,1000000,2500000 HP1.jmatrix HP1.hic hg38

java -Xmx120g -jar juicer_tools_1.22.01.jar pre --threads 40 -r 5000,10000,25000,50000,100000,250000,500000,1000000,2500000 HP2.jmatrix HP2.hic hg38

for res in 10000 50000 100000 250000 500000;do

hicConvertFormat --matrices HP1.hic --inputFormat hic --outFileName HP1.$res.cool --resolutions $res --outputFormat cool

hicConvertFormat --matrices HP2.hic --inputFormat hic --outFileName HP2.$res.cool --resolutions $res --outputFormat cool

cooler balance --cis-only HP1.$res.cool

cooler balance --cis-only HP2.$res.cool

done

With these cool files we can examine the Pore-C diploid contacts from different points. We first compute the compartment eigenvector scores with 100 kb resolution (*cis.vecs.tsv):

for res in 10000 50000 100000 250000 500000;do

cooltools eigs-cis --bigwig -o HP1.$res HP1.$res.cool

cooltools eigs-cis --bigwig -o HP2.$res HP2.$res.cool

done

Compute the TAD insulation scores (*insulation.tsv). We compute insulation scores with cooltools under 50 kb resolution and 2, 5, 10 sliding windows.

cooltools insulation -o HP1.insulation.tsv --window-pixels --append-raw-scores --bigwig HP1.10000.cool 1 2 5 10

cooltools insulation -o HP2.insulation.tsv --window-pixels --append-raw-scores --bigwig HP2.10000.cool 1 2 5 10

Compute the Loop apa scores (enhancement.txt). We perform structural loops aggregate enrichment analysis on based GSE63525 to show that Pore-C diploid contacts matrices can successfully detect loops using juicer tools apa.

java -Xmx20g -jar juicer_tools_1.22.01.jar apa --threads 15 -r 10000 -w 10 -u HP1.hic GSE63525_GM12878_primary+replicate_HiCCUPS_looplist.txt loops_HP1/

java -Xmx20g -jar juicer_tools_1.22.01.jar apa --threads 15 -r 10000 -w 10 -u HP2.hic GSE63525_GM12878_primary+replicate_HiCCUPS_looplist.txt loops_HP2/

### Supplementary Note 8: Assessment of Pore-C diploid genome

Detecting refined 3D structures requires higher resolution and data coverage. Generally, with smaller bin size and larger number of contacts we can obtain a clearer 3D genome. The library preparation stage of Pore-C will inevitably introduce noises such as random-ligation and self-ligation. Besides, in the fragment haplo-tagging stage of Dip3D, incorrectly tagged contacts will ruin the consistency and richness of contacts. Hence it is reasonable to assess the quality of a diploid genome from uniformity of Pore-C fragment coverage and consistency.

According to Rao et al., a diploid 3D genome is said to achieve solution of $k$, if over 80% of its bins (each with equal size of $k$) have interacting counts greater than 1000. We use QuASAR-QC to evaluate the quality of a single contact matrix and use HiCRep to evaluate the correlation between maternal and paternal contact matrices. All these evaluations are integrated into 3DChromatin_ReplicateQC (v1.0.1).

We first computed the interaction frequencies with different bin size (*ginteractions.tsv):

for res in 10000 50000 100000 250000 500000;do

hicConvertFormat --matrices HP1.$res.cool --inputFormat cool --outFileName HP1.$res.ginteractions --outputFormat ginteractions --load_raw_values

hicConvertFormat --matrices HP2.$res.cool --inputFormat cool --outFileName HP2.$res.ginteractions --outputFormat ginteractions --load_raw_values

done

And perform the evaluations

3DChromatin_ReplicateQC run_all --metadata_samples metadata.samples --metadata_pairs metadata.pairs --bins bin.txt.gz --outdir 3D_RepQC_out/

Benefit from the longer fragment length and advantages of third generation sequencing, Pore-C successfully reconstructed diploid 3D structures on regions that are inaccessible to Hi-C, such as GC-contents, segmental duplications, and regions with low mappability. These difficulties can be partially resolved with ultra-high sequencing depth of Hi-C.

The difficulty regions used in this study come from GIAB. Take GRCh38_gc15_slop50.bed as an example, we conducted coverage stats with following commands:

samtools cat -@ 40 */tagged-*.bam | samtools sort -@ 40 o tagged.bam

samtools index tagged.bam

samtools view -d HP tagged.bam > tagged-only.bam

samtools index tagged-only.bam

total_bases=`awk -F'\t' '{print $3-$2}' GRCh38_gc15_slop50.bed |awk '{sum+=$1} END {print sum}'`

cover_bases=`samtools depth -@ 40 -b GRCh38_gc15_slop50.bed tagged-only.bam |wc -l`

### Supplementary Note 9: Reproduction of canonical haplotype-specific 3D structures

For comparison with Hi-C, we extracted diploid contact matrices from the high order contacts of Dip3D outputs based on 190× HG001 Pore-C data. We first compared the diploid 3D structure of HG001 by Dip3D to that previously published based on Hi-C (~878×)^2^. Rao et al. (2014) reported notable haplotype-specific structures associated with X chromosome inactivation (XCI) in human females^2,3^. The diploid 3D structures of chrX was then compared at levels of A/B compartment and TAD (Supplementary Figure 2A,B). In respect of the two levels, Pore-C and Hi-C based 3D structures both had higher correlations between homologous autosomes (Supplementary Figure 2C) than between the two X-chromosomes. The Pore-C matrix successfully identified the superdomain (Supplementary Figure 2D-F) located at the ~115 Mb coordinate of the deactivated paternal X-chromosome, as well as the superloops (Supplementary Figure 2G) formed by two tandem repeat regions DXZ4 and FIRRE, which interact with CTCF structural protein and are involved in the remodeling process of the inactivated X chromosome^4^.

Another example of haplotype-specific 3D structure is related to the genetic imprinting, such as the *H19* and *Igf2* genes on the autosome chr11 that competitively bind to the nearby *H19*/*Igf2* Distal Anchor Domain (HIDAD) locus^2,5^. *H19*-HIDAD interaction dominates in the maternal-origin chromosome, while the two interaction types (*H19*-HIDAD and *Igf2*-HIDAD) are more evenly present in the paternal-origin chromosome^5^. Using the Dip3D diploid contact matrix, we calculated the frequencies of *H19*/*Igf2*-HIDAD interactions in the maternal- and paternal-chromosomes of NA12878. We found that H19-HIDAD interactions within the maternal-haplotype occurred twice as frequently as within the paternal-haplotype (Supplementary Figure 2G).
